# pA regulator: a system for controlling mammalian gene expression via the modulation of polyA signal cleavage

**DOI:** 10.1101/2023.01.27.525935

**Authors:** Liming Luo, Jocelyn Duen-Ya Jea, Yan Wang, Pei-Wen Chao, Laising Yen

**Affiliations:** Department of Pathology & Immunology, Department of Molecular and Cellular Biology, Dan L. Duncan Cancer Center, Baylor College of Medicine, Houston, TX 77030, USA

## Abstract

The ability to control the expression of a therapeutic gene or a transgene in mammalian cells is crucial for safe and efficacious gene and cell therapy, as well as for elucidating the function of a specific gene product. Yet current mammalian gene regulation systems either evoke harmful immune responses in hosts or lack the required regulatory efficiency. Here we describe a highly responsive RNA-based molecular switch, the pA regulator, that harnesses the power of polyA signal cleavage within the 5’ UTR to control mammalian gene expression. The pA regulator is governed by a ‘dual mechanism’ to ensure maximal control of gene expression: (1) aptamer clamping of polyA signal via drug binding and (2) drug-induced alternative splicing that removes the polyA signal. The metholology achieves an induction efficiency up to 900-fold with an EC_50_ of 0.5μg/ml Tetracycline, a drug concentration that falls well within the FDA-approved dose range. The pA regulator circumvents the immune responses that plague other systems by eliminating the use of a regulatory foreign protein and the need to change transgene coding sequences. Furthermore, it is not dependent on any specific promoter, therefore the system is simple to implement in a single non-viral or viral vector. In a mouse study using AAV-mediated gene transfer, we showed that the pA regulator controlled transgene expression in a “dose-dependent’ and “reversible” manner and exhibited long-term stability *in vivo*, in which both features are crucial for effective therapeutics. The pA regulator is the first non-immunogenic system that demonstrates an EC_50_ at a drug concentration approved by FDA, making it a clinically relevant gene regulation system that could open a new window of opportunity in clinical applications as well as biological studies.

## Introduction

The ability to control the expression of exogenous therapeutic genes to appropriate levels within the ‘therapeutic window’ is important for the safety and efficacy of gene and cell therapy as over-expression could cause severe toxicity and even death^1, 2^. With the complexity of diseases increasingly understood today, it is clear that effective therapies catered to the needs of individuals will require fine-adjustment or time-varying expression according to disease stages or severity^3^. These considerations call for the development of a safe and efficient gene regulation system to control the levels of therapeutic gene products after gene delivery. However, current gene transfer methods used in gene and cell therapy, such as Adeno-Associated Virus (AAV), have intrinsic difficulties in executing the ‘conditional’ and ‘reversible’ control of genes that often need additional calibrations after delivery^3, 4^. While the commonly used Tet-on system proved to be a powerful gene regulation tool in biological studies^5^, it is well-documented that its use of foreign regulatory ‘transactivator’ protein can trigger immune responses in nonhuman primates^6–8^. As a result, there are no approved clinical applications based on the Tet-on system. In addition, the host’s intolerance to foreign protein epitopes can be extremely discerning, such that a therapeutic protein with a single amino acid difference to the native host human protein is sufficient to evoke a transgene-specific immune response in patient^9^. In fact, immune responses against foreign protein epitopes remains a key hurdle to successful gene and cell therapy as it causes elimination of gene expression and destruction of transduced host cells^6–8, 10, 11^. These concerns highlight the importance of using a non-immunogenic gene regulation system for safe and efficacious gene and cell therapy.

The need for a non-immunogenic gene regulation system prompted the development of numerous RNA-based systems with the objective of achieving gene regulation in mammalian cells without the use of any foreign protein epitopes^12–19^. These RNA-based systems have invariably exhibited fundamental limitations in regulatory efficiency such as the use of toxic ligand, narrow dynamic ranges, high leakage expression, and above all, the requirement of high ligand concentrations for induction^4, 17^. In fact, published RNA-based systems commonly neglect to state their EC_50_ (the ligand concentration required for half-maximal induction). Yet, available results in the literature^4, 17^ suggest that the EC_50_ exhibited by these RNA-based systems in mammalian cells is likely at least a magnitude higher than what is useful for both biological studies and clinical applications. For this reason, none of these systems have moved beyond the proof-of-concept stage for any practical application to date. Recently, an RNA-based system designated ‘X-on switch’ that utilizes drug-induced alternative splicing to achieve gene regulation in mammalian cells was proposed as a possible solution^20^. Such a system however adds a stretch of unrelated amino acids to the transgene coding sequence, thus promoting a transgene-specific immune response that RNA-based systems seek to avoid.

In this manuscript, we describe the ‘pA regulator’, a mammalian gene regulation system that fills this critical gap by overcoming the key issues associated with existing gene regulation systems. The success of this RNA-based system depends on polyA signal (PAS) cleavage to modulate gene expression without the use of a foreign regulatory protein. Also, it allows for the expression of any intact protein as a transgene product without changing the coding sequence. These features circumvent the transgene-specific immune responses that have plagued the use of other systems. Moreover, the pA regulator provides a regulatable range up to 900-fold while limiting the basal leakage expression to 0.1%. Most importantly, the system exhibits a switching efficiency with an EC_50_ of 0.5μg/ml inducer molecule (tetracycline in this case). As such, it is the only ‘non-immunogenic’ gene regulation system to date that demonstrates an EC_50_ at a drug concentration approved by the FDA. These advances make the pA regulator a safe and efficient regulation system that could be useful for broad clinical applications as well as diverse biological studies.

## Modulation of polyA signal cleavage via Tc-binding aptamer

The central strategy for controlling gene expression through modulation of PAS cleavage is shown in Fig. 1A. The approach is critically dependent on two principles. First, a synthetic PAS is placed in the 5’UTR of a transgene to generate highly efficient cleavage of the corresponding mRNA molecule. This cleavage leads to the destruction of the coding region of the mRNA and therefore loss of transgene expression. Second, a small molecule ligand inhibits the cleavage via binding to an incorporated RNA aptamer sequence surrounding the PAS. When engineered properly, ligand binding strengthens the RNA aptamer structure and efficiently blocks the cleavage of the synthetic PAS, resulting in preservation of the intact mRNA, therefore inducing transgene expression.

**Figure 1.**
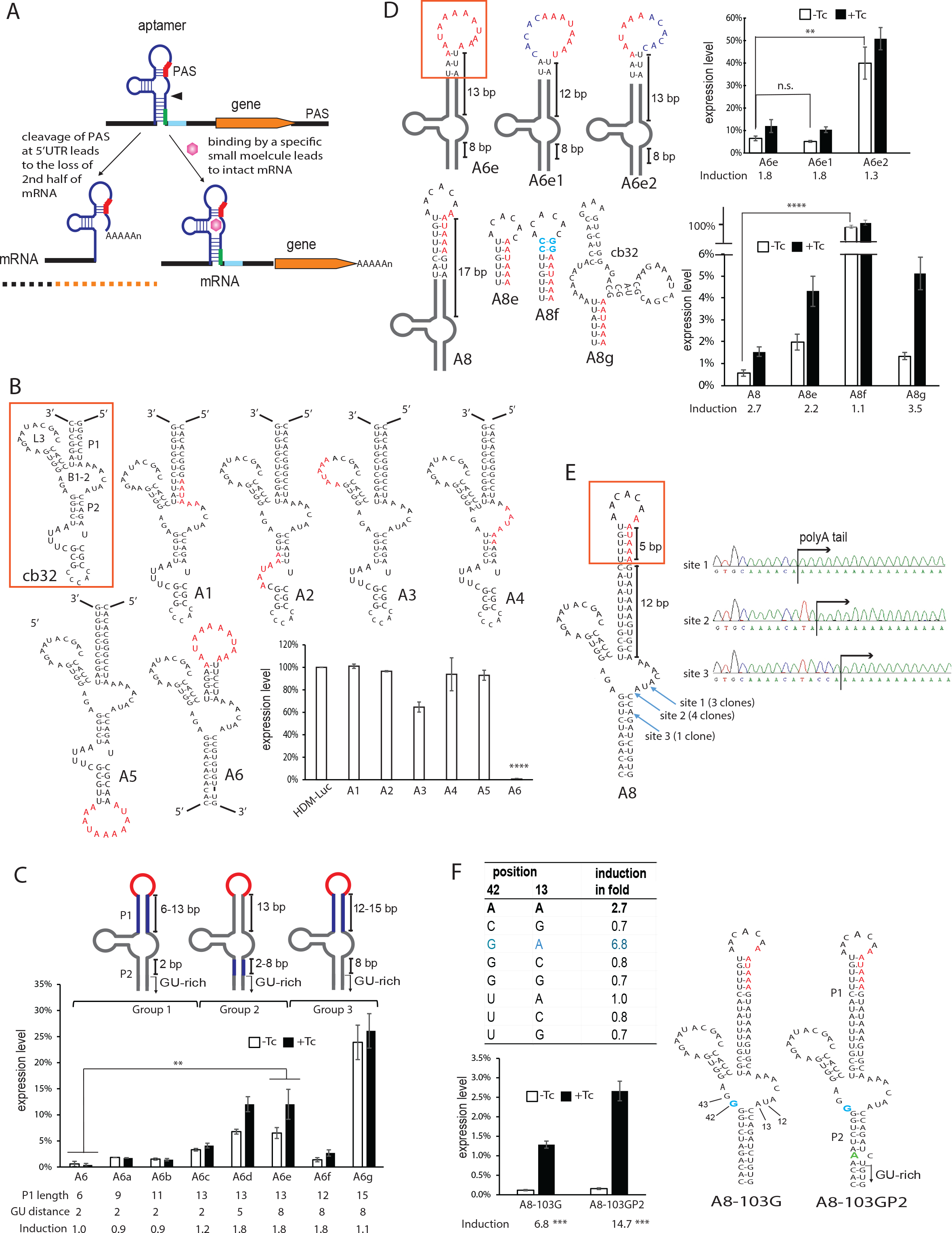
Modulation of PAS-mediated cleavage via Tc-binding aptamer. **A**. Strategy for controlling gene expression via PAS cleavage. A synthetic PAS (red) is inserted in the 5’UTR of a transgene to generate efficient cleavage of its mRNA molecule. In the absence of ligand, the cleavage leads to the destruction of the coding region of the mRNA and therefore loss of transgene expression. Binding of the ligand to the aptamer inhibits the cleavage, resulting in preservation of the intact mRNA, therefore induced transgene expression. A GU-rich region (green) and a G-rich motif MAZ (blue) are added to augment polyadenylation. The entire strategy is based on a single molecule of RNA, and all engineered RNA motifs are cis-acting. The arrowhead points to the cleavage site. **B**. Probing locations within aptamer that allow synthetic PAS to function efficiently. The cb32 aptamer (boxed) is used to host the synthetic PAS (red letters). The L3 loop and the B1-2 bulge constitute the Tc-binding core of cb32 aptamer. PAS was tested in locations where minimum changes of the aptamer sequence are needed to create “AAUAAA”, and/or where PAS can be placed partially in the single-stranded region. The relative luciferase activity was measured in the absence of Tc. The activity of the parental plasmid (HDM-Luc) was used as the relative 100%. Lower luciferase expression in the absence of Tc indicates higher PAS cleavage efficiency. p<0.01 for expression level, HDM-Luc versus A6. **C**. The effects of P1 stem length and the position of GU-rich region on induction. Constructs were tested for luciferase expression in the absence (white bars) or presence of 15 μg/mL Tc (black bars). Group 1: Constructs with P1 stem length varied from 6 to 13bp. Group 2: Constructs with the distance of GU-rich region to the B1-2 bulge varied from 2 to 8 bp. Group 3: Constructs with P1 length varied from 12 to 15 bp while GU-rich region distance fixed at 8bp. Red: PAS. Blue: area of modification. The regulation efficiency of each construct was measured as ‘induction in fold’ which is a ratio of luciferase reporter signal in the presence vs. absence of Tc. Error bars represent standard deviations. p<0.01 for induction in fold, A6 versus A6e. **D**. The effects of copy number and clamping of PAS. PAS in red letters. A6e1: the first PAS mutated; A6e2: the second PAS mutated. A8: PAS partially embedded in P1 stem. A8e: PAS completely embedded in the P1. A8f: PAS clamped by additional two C-G pairs (blue). A8g: PAS clamped by an additional cb32. Statistics compare the leakage background expression. **E**. Identification of PAS cleavage sites by RT-PCR. Sanger sequencing (right) of the RT-PCR products revealed three cleavage sites (blue arrows) where the poly-A tail was added to the RNA. Sites 1 and 2 located in bulge B1-2 appeared more frequently than site 3. **F**. Mutational analyses on positions A42/A13 based on the A8 construct. The table shows that an A42G mutation (blue letters) more than doubled the induction (A8-103G). A spontaneous G to A mutation (green) in P2 stem also more than doubled the induction (A8-103GP2). The schematics on right indicates the locations of those beneficial mutations. Statistics compare induction in fold, p<0.001 for A8 versus A8-103G, p<0.001 for A8-103G versus A8-103GP2. Data shown are average (mean ± SD) of representative experiment performed independently at least three times, each with triplicated cell cultures. Significance was determined by two-tailed Student’s t-test; n.s.: not significant, **: p<0.01, ***: p<0.001, ****: p<0.0001.

## Identification of locations within the aptamer that allow PAS to function efficiently at the 5’ UTR

The success of this approach relies on the coupling of PAS and the aptamer to function as an integrated unit at the 5’UTR. Although PAS-mediated cleavage is a strong mRNA processing mechanism universally present in all mammalian cells, PAS normally locates at the 3’ UTR but not 5’UTR of mRNA. Moreover, strong surrounding RNA structures are known to inhibit PAS cleavage ^21–23^. Therefore, our first step was to determine configurations where a synthetic PAS embedded in a structured aptamer can remain highly active within the 5’UTR of a mammalian mRNA. We chose the well-characterized ‘cb32’ aptamer ^24, 25, 26^ that binds tetracycline (Tc) as the ‘sensing’ structure to host and control the synthetic PAS. Tc is an FDA-approved drug that is orally bioavailable and has been widely used in clinics for decades^27, 28^.

As shown in Fig. 1B, using the cb32 aptamer as the host sequence, we introduced the synthetic PAS in several strategic locations where minimum substitutions of the aptamer sequence are needed to create the PAS sequence “AAUAAA”, and/or where PAS can be placed at least partially in the single-stranded region to minimize structural obstruction. Furthermore, we added two downstream elements^29^ to enhance the function of synthetic PAS: (1) a GU-rich region^30^ and (2) a G-rich motif called MAZ^31–33^ known to augment polyadenylation (Fig. S1). These motifs were inserted after the aptamer in the 5’UTR of a standard mammalian expression vector which encodes a luciferase reporter gene (Fig. 1A)^12^. To measure the cleavage efficiency of the inserted synthetic PAS, human 293T cells were transiently transfected with the respective plasmids, and one day later, the luciferase activity was measured in the absence of Tc. If the synthetic PAS leads to efficient cleavage of the reporter mRNA, then a loss of luciferase expression is expected. As shown in Fig. 1B, inserting PAS in or near the binding core of the aptamer led to little or no loss in luciferase expression (Fig. 1B, construct A1 to A4). In contrast, inserting two copies of PAS as an open loop at the distal end of P1 stem (Fig. 1B, construct A6) greatly reduced luciferase expression to less than 1% of that of the parental vector (HDM-Luc). However, placing the same loop at the end of P2 stem failed to reduce gene expression (Fig. 1B, construct A5). Together, the results suggest that PAS is capable of functioning efficiently when embedded in the Tc aptamer, albeit the insertion locations need to be empirically determined. Furthermore, there is no ‘intrinsic’ cellular inhibition of PAS inserted at the 5’ UTR of mRNA, an exotic location where PAS never localizes in normal mammalian transcripts.

Based on its apparent high efficiency of PAS-mediated cleavage, A6 was chosen as the base configuration for developing a molecular switch that can be controlled by Tc. To quantify the induced gene expression in response to Tc, we measured the ‘induction in fold’ as a ratio of luciferase reporter signal in the presence vs. absence of Tc. As shown in Fig 1C, A6 exhibited no appreciable increased expression in response to Tc (fold=1) in cells, indicating that Tc binding produces no effect on PAS that is located at the distal end of the P1 stem. To identify a configuration that allows Tc binding to exert influence on the PAS, we varied the length of the P1 stem from 6bp to 9, 11, and 13bp respectively (Fig. 1C, group 1, A6-A6c; detailed sequences in Fig. S2). This led to a moderate but measurable induction of 1.2-fold when P1 is increased to 13bp. Next, we probed the effect of polyA cleavage sites within the aptamer by altering the distance of the GU-rich region to PAS, as this distance is known to determine the polyA cleavage locations that lie in between^30^. As shown in Fig. 1C, increasing the distance of the GU-rich region relative to the aptamer core (therefore the absolute distance from GU-rich region to PAS) from 2bp to 5 or 8bp further improved the induction to 1.8-fold (Fig.1C, group 2, A6c-A6e, and Fig. S2). With this distance maintained at 8bp, shortening the P1 length from 13bp to 12bp did not change the induction while lengthening it to 15bp reduced the induction to 1.1-fold due to the higher basal background expression. These results indicate that a distance of 13bp from PAS to the core is near optimal. In addition, subtle changes in the lengths of P1 and P2 greatly affects the switching performance, as they affect not only the overall stability of the RNA structure where the PAS is embedded but also the accessibility of PAS cleavage locations within the aptamer structure.

Because all constructs in the A6 series contain two copies of the PAS in the loop, to determine which PAS contributes more to the cleavage, we inactivated each unit individually by replacing the sequence ‘AAUAAA’ with ‘ACACAC’. Inactivating the first PAS in A6e exerted little difference in both the basal expression and the fold induction. In contrast, inactivating the second PAS led to a dramatic increase in basal expression and reduced the induction (A6e2 vs. A6e1, Fig.1D and S3). Therefore, the second PAS, but not the first, is responsible for efficient cleavage, and a single PAS is sufficient to maintain the performance of A6e.

To exert a greater influence of Tc binding on the synthetic PAS, we incorporated part of the single PAS into the P1 stem by adding nucleotides to base pair with the PAS, while the distance of the PAS to the aptamer core remained unchanged. This increased induction from 1.8 to 2.7-fold (Fig. 1D, A8). However, burying the PAS in the P1 stem (A8e) resulted in higher basal background expression therefore a lower fold-induction as compared to A8. Most importantly, clamping the PAS by two additional C-G pairs (A8f) restored nearly 100% luciferase expression in the absence of Tc. This result suggests that, in theory, it is possible to achieve ‘very efficient’ control of PAS cleavage through modulation of structural stability near the PAS. Consistent with this idea, adding another Tc aptamer next to the PAS without elaborate optimizations quickly improved induction from 2.7 to 3.5-fold (A8g) albeit the improvement is moderate.

To identify the PAS cleavage locations within the aptamer, we amplified the polyadenylated RNA expressed from construct A8 using a dT oligo paired with a 5’ primer specific to the aptamer. Sanger sequencing of the cloned RT-PCR products identified three cleavage sites where the polyA tail is added to the aptamer RNA sequence by polyadenylation (Fig.1E). All of the identified cleavage sites, which lie between the synthetic PAS and GU-rich region, were located either inside (site 1 and 2, with higher frequencies) or close to the Tc binding core (site 3). This suggests that Tc binding, in addition to stabilizing the overall RNA structure, also directly blocks the cleavage sites, and both contribute to the overall inhibition of PAS cleavage.

Most of the bases in the binding core of cb32 aptamer^24,25^ have been optimized by mutational analyses^34^. One exception is position A42 (Fig. 1F), which is known to contact position A13 by hydrogen bonds^25^, but their mutational effects have not been reported. We mutated both positions and found that the A42G substitution (A8-103G) more than doubled the induction from 2.7-fold to 6.8-fold (Fig. 1F, the table), while other mutations in these positions abolished induction. The surprising results suggest that the A42G substitution, which is present in an earlier variant of Tc aptamer called ‘cb12’^26^, enhances the switching response of aptamer to Tc. Moreover, we accidentally isolated a spontaneous G to A mutation in the P2 stem during cloning, which doubled the induction from 6.8-fold to 14.7-fold (Fig. 1F, A8-103G vs. A8-103GP2). This mutation is unlikely to have a direct influence on the PAS and the binding core, as its location is away from both. However, the resulting change from a G-C pair to an unpaired A/C in the P2 stem may have modulated the stability near the GU-rich region, a motif important for the efficiency of PAS cleavage.

## The Y-shaped structure converges the binding effects from three aptamers

Having developed the A8 configuration (and its improved version A8-103GP2) that enables the synthetic PAS to function sufficiently, we then turned our attention to the important observation that the activity of PAS can be abolished by as little as two additional C-G pairs (Fig. 1D, A8f). This implies that replacing the clamping force of two C-G pairs with an equivalent force generated by Tc binding could lead to substantially improved induction. To exert such conditional suppression on the PAS, we used A8-103GP2 that contains the A42G mutation (designated as aptamer “A”) as the scaffold and engineered two additional aptamers (“B” and “C”) on top of the PAS. The resulting ‘Y-shaped’ structure (Fig. 2A) contains three comparable arms (Arms 1, 2, 3 respectively), each transmitting the effect from one aptamer toward the central loop that constitutes a 3-way junction. In this new configuration, part of the PAS sequence is in the central loop, which is reminiscent of the loop of A8-103GP2 that allows efficient PAS cleavage in the absence of Tc. Upon Tc binding, the Y-shaped structure is expected to channel the clamping force from each aptamer and converge the effects on the 3-way junction where the PAS resides.

**Figure 2.**
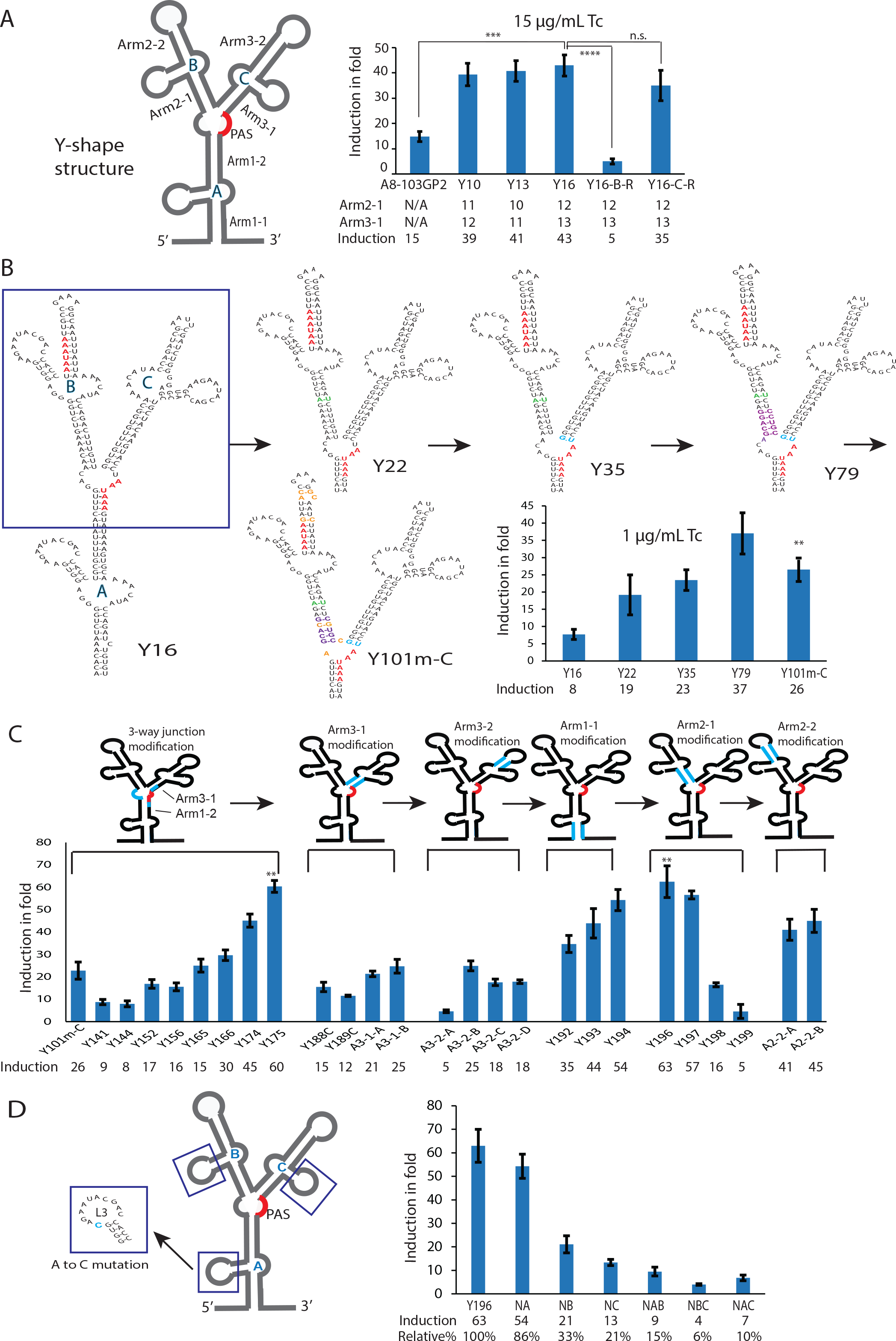
The Y-shape structure converges the binding forces from three aptamers. **A**. Left: Schematics of the Y-shape structure. Aptamer A, B, C are connected to the 3-way junction through their P1, P2, P1 stem respectively. The synthetic PAS is in the central 3-way junction marked in red. We labelled each arm and aptamer according to their order of appearance in the mRNA sequence. Right: Regulation efficiency of various Y-shape constructs. Y10, Y13 and Y16 have Arm2-1 and Arm3-1 varied from 10 to 13 bp. Y16-B-R has aptamer B orientation reversed relative to that of Y16. Y16-C-R has aptamer C orientation reversed relative to that of Y16. N/A: not applicable. **B**. The modification of various parts of Y-shape structure that led to noticeable improvements in induction. The areas of modification are indicated by colors and the resulting regulation efficiency are shown in the bar chart. Y22 lengthens Arm2-1 by an A-U pair (green). Y35 has a U to G change that lengthened Arm3-1 (blue). Y79 has changes from A-U to G-C pairs (purple) that further stabilizing Arm2-1. Y101m-C contains several specific modifications (orange) including changes in the central loop, the removal of a potential 3’ss ‘AG’ in Arm2-1, and the elimination of a potential PAS in Arm2-2. p<0.01, Y16 versus Y101m-C. **C**. Effects of synthetic PAS locations in the 3-way junction and the fine-turning of arm lengths. The areas of modification are indicated by blue and the resulting regulation efficiency are shown in the bar chart. Modifications of 3-way junction are based on Y101m-C as the parental. Modifications of Arm 3-1, Arm3-2, Arm1-1, and Arm2-1 are based on Y175 as the parental. Modifications of Arm2-2 are based on Y196 as the parental. Details of modifications are shown in Fig. S5 to S8. p<0.01, Y101m-C versus Y175; p<0.01, Y101m-C versus Y196. **D**. Asymmetric contribution of each aptamer to the overall induction. Left: An “A” to “C” mutation (blue) in the binding pocket (squares), which eliminates the binding of Tc, is used to disable aptamer A, B, or C individually (NA, NB, and NC respectively), or to disable two aptamers at once (NAB, NBC and NBC respectively). Right: Regulation efficiency. The results indicate that aptamer C is the most important among the three, followed by aptamer B and then A. Statistics compare induction in fold. Data shown are average (mean ± SD) of representative experiment performed independently at least three times, each with triplicated cell cultures. Significance was determined by two-tailed Student’s t-test; n.s.: not significant, **: p<0.01, ***: p<0.001, ****: p<0.0001.

To quickly probe whether the designed Y-shaped structure functions as predicted, and to determine the ideal arm lengths that channel the force from the aptamers B and C to the central loop, we designed a series of Y-shaped constructs with Arm2-1 and Arm3-1 that varies from 10 to 13 bp (Fig. S4, top panel). As shown in Fig. 2A, all these designs (constructs Y10, Y13, and Y16) nearly tripled the induction, indicating the superb effectiveness of the Y-shaped configuration. Among them, Y16 (arm lengths=12bp/13bp) performed the best, giving a 43-fold induction as compared to 14.7-fold of the parental A8-103GP2. Furthermore, the orientation of aptamer A, B, C in the Y-shaped structure was designed to facilitate correct individual aptamer folding and to minimize incorrect folding that can be formed among the three similar aptamers. Indeed, inverting the orientation of aptamer B in Y16 (Y16-B-R), which permits misfolding between aptamer A and B, dramatically reduced the induction from 43-fold to 5-fold, while inverting the orientation of aptamer C (Y16-C-R) slightly reduced the induction to 35-fold (Fig. 2A and Fig. S4). Thus, the specific orientation of the aptamers is critical for the Y-shaped structure to function properly.

To optimize the configuration of Y16, a series of modifications were made to probe the effects of various parts of the Y-shaped structure. These constructs were then tested at 1μg/ml Tc (equivalent to approximately 2μM Tc), as opposed to the 15μg/ml Tc used previously, so that specific modifications that allow efficient induction at 1μg/ml Tc (a serum concentration can be reached by taking FDA-approved Tc dosage) could be identified. Fig. 2B highlights the changes that led to noticeable improvements. Lengthening Arm2-1 by an A-U pair (green in Y22), a U to G change that lengthened Arm3-1 (blue in Y35), and further stabilizing Arm2-1 by converting A-U to G-C pair (purple in Y79), progressively improved the induction from 8-fold to 19-, 23- and 37-fold respectively (Y16 vs. Y22, Y35, and Y79) at 1μg/ml Tc. The construct Y101m-C contains several specific modifications (orange), notably those involving the central loop, the removal of a potential 3’ splice site ‘AG’ in Arm2-1, and the elimination of a potential PAS in Arm2-2. While these modifications moderately decreased the performance, they simplify the engineering principles by removing unwanted cryptic elements. For this reason, Y101m-C was chosen as the basis for subsequent studies. Investigating the effects of PAS location within the 3-way junction showed that embedding it entirely in Arm1-2 led to markedly reduced the induction (Fig. 2C and Fig. S5, Y141/Y144 vs. Y101m-C). Embedding the PAS as part of Arm1-2 and Arm3-1 also reduced the induction (Y152/Y156 vs. Y101m-C). However, when the PAS is partially open in the central loop, it yielded generally improved performance (Y165/Y166/Y174/Y175 vs. Y101m-C). Among these, Y175 struck an optimal balance with 4 nts of PAS in the central loop and 2 nts embedded in Arm1-2, which led to the highest induction of 60-fold at 1μg/ml Tc.

In contrast, decreasing or increasing the stability of Arm3 by altering its length in Y175 led to a much-reduced induction (Fig 2C, Arm3-1 and Arm3-2 modification). Similarly, decreasing or increasing the stability of Arm1 in Y175 also reduced the induction (Fig 2C, Arm1-1 modification). These results indicate that the stabilities of Arm3 and Arm1 of Y175 are nearly optimal. Altering the stability of Arm2 led to variable effects (Fig 2C, Arm2-1 and Arm2-2 modification). Among them, Y196 showed slightly improved performance of 63-fold as compared to its parental Y175. Detailed sequences of these modifications are shown in Fig. S5 to Fig. S8.

Having optimized the location of PAS and stability of the three arms of the Y-shaped structure, we then asked whether the three aptamers contributed equally to the final induction. To answer this question, we disabled each aptamer individually by an ‘A to C’ mutation in the binding pocket known to eliminate Tc binding^34^. Disabling aptamer A, B, or C (NA, NB, and NC respectively) resulted in the loss of induction by 13%, 67%, and 79% respectively as compared to the parental Y196 (Fig. 2D). Disabling two aptamers at once (NAB, NBC, and NAC) led to further loss of induction (by 86%, 94%, and 89% respectively). These results indicates that while every aptamer in the Y-shaped structure contributes to the final induction, the contribution is asymmetrical, with aptamer C being the most influential followed by aptamer B, and then A. Furthermore, induction does not require simultaneous Tc binding to all three aptamers, as binding to any one aptamer alone is sufficient to induce various levels of gene expression.

## Harnessing the power of alternative splicing as a second mechanism for controlling PAS

The continuous and labor-intensive step-by-step improvement described above led to Y196 with a respectable induction of 63-fold, which already surpasses the performance of previous published RNA-based switches at 1μg/ml Tc. However, many practical applications would require a larger dynamic range of gene regulation. Yet, a switching mechanism that solely relies on the clamping effect of Tc binding onto the synthetic PAS may have already reached its practical limitation in the configuration of Y196, which is the product of extensive optimizations. To achieve the next level of effectiveness and dynamic range, we instilled a second parallel mechanism to control the synthetic PAS: namely drug-inducible alternative splicing. Past reports indicated that binding of cellular factors to a specific G-quad structure influences the choices of alternative 3’ splice sites (3’SS)^35, 36^, presumably by altering the local RNA structural stability. As shown in Fig. 3A, all our constructs contain a built-in IVS2 intron in the 5’UTR that enhances reporter gene expression, as well as a G-quad motif called MAZ that enhances the activity of synthetic PAS. Based on this existing configuration, we implemented a strategy of drug-inducible alternative splicing by adding an alternative 3’SS downstream of the G-quad. The rationale is illustrated in Fig. 3A. In the absence of Tc, the IVS2 intron is correctly spliced, and the cleavage of the PAS leads to the degradation of the mRNA, disrupting gene expression. Tc binding is predicted to strengthen the Y-shaped structure located near the MAZ G-quad; when engineered properly, this would lead to induced alternative splicing to the 3’SS downstream of the G-quad. The resulting alternative splicing is designed to serve two critical purposes. First, it provides a second independent mechanism to preserve the mRNA by splicing out the PAS entirely. Second, it removes the bulky Y-shaped and the G-quad structure from the 5’UTR that may hinder protein translation^37, 38^.

**Figure 3.**
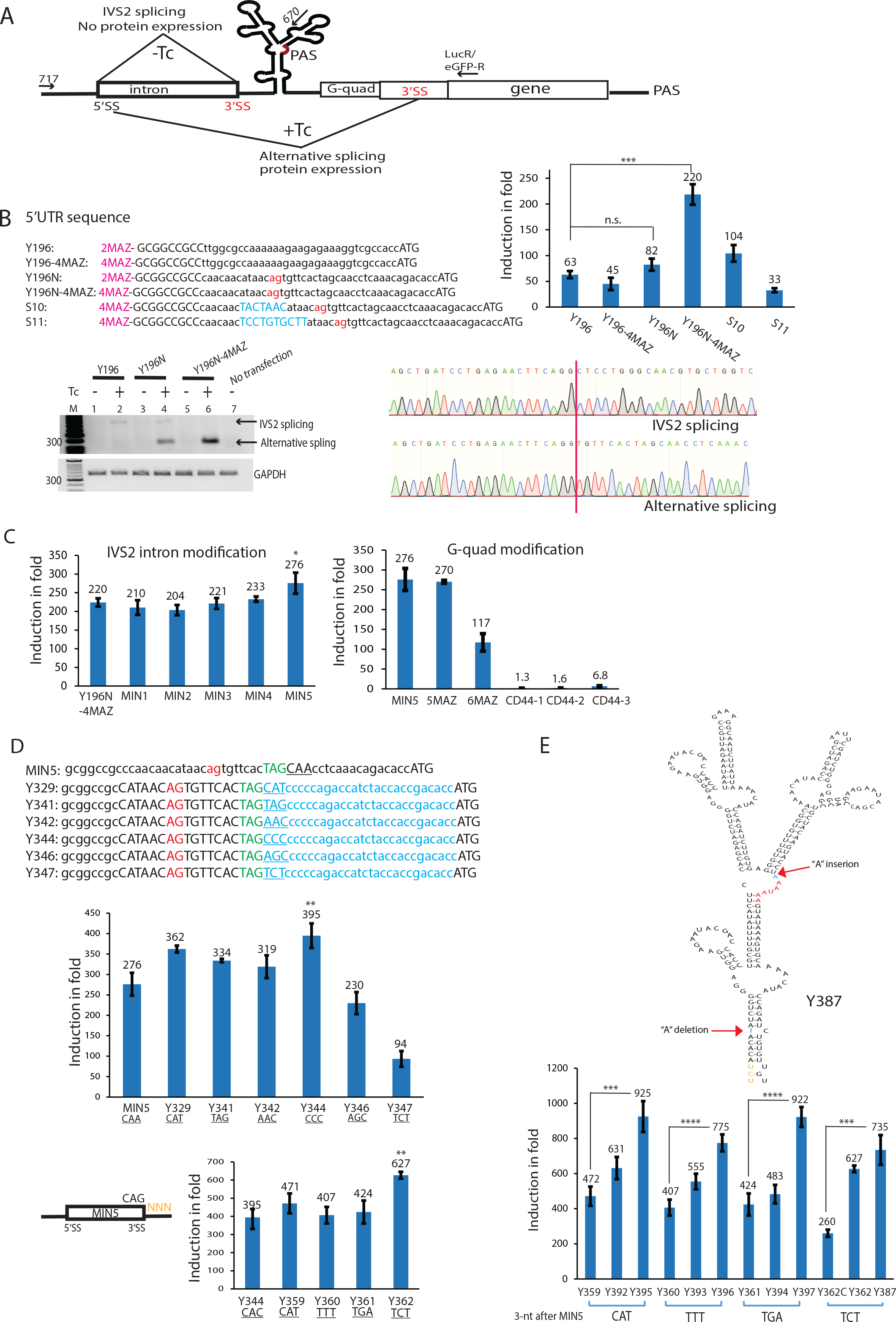
Harnessing the power of alternative splicing as a second mechanism for controlling PAS. **A**. Strategy for combining PAS cleavage and alternative splicing to control gene expression. In the absence of Tc, the pre-mRNA undergoes normal IVS2 splicing and the cleavage of synthetic PAS leads to the destruction of transgene mRNA and therefore the loss of protein expression. Binding of Tc to the aptamers in the Y-shape structure inhibits the PAS cleavage, and at the same time, induces alternative splicing. The alternative splicing removes the PAS, the Y-shape and the G-quad structure from the pre-mRNA, which results in the preservation of transgene mRNA, therefore the induced protein expression. **B**. Engineering alternative splicing by modifying the 5’UTR with 3’SS and G-quad sequence. Top left: The various 5’UTR sequences combined with two or four copies of MAZ. Top right: Their induction efficiency. Bottom left: RT-PCR reveals the mechanism of drug-induced alternative splicing. Primer locations are marked by arrows in **A**. Primer pair 717/LucR detects both the IVS2-spliced and alternatively spliced luciferase mRNA. GAPDH is used as loading control. In the absence of Tc, IVS2-spliced RNA is degraded by PAS cleavage (lane 1, 3 and 5). The presence of Tc induces alternative splicing in both Y196N and Y196N-4MAZ (lane 4 and 6, band of 193 bp), but not in Y196 (lane 2). Tc-induced alternative splicing is far more pronounced in Y196N-4MAZ than in Y196N (lane 6 vs. 4). Bottom right: Sanger sequencing confirmed the expected IVS2 splice junction in the absence of Tc, and the alternative splice junction in the presence of Tc. The red vertical line marks the splice junctions. The identified alternative 3’SS “AG” is marked in red in the sequences in the Top left panel. Branch point in S10 and polypyrimidine tract in S11 are marked in blue. **C**. Effects of IVS2 intron size and G-quad sequence. Left: The effect of shortening IVS2. IVS2 intron of Y196N-4MAZ has a size of 476 nts. MINI1 to MINI4 introns have sizes of 180, 160, 140, 120 nts respectively. MIN5 further mutates an ‘ATG’ present in the MIN4 intron. Right: The effect of G-quad sequence and copy numbers. Constructs with four (MIN5), five (5MAZ), and six (6MAZ) copies of MAZ, and one (CD44-1), two (CD44-2), and four (CD44-3) copies of CD44 G-quad are compared. p<0.05, Y196N-4MAZ versus MIN5. **D**. Modifications of 5’UTR sequence to optimize the balance between normal splicing and drug-induced alternative splicing. Top panel: Various 5’UTR with unstructured sequences (blue) inserted before the start codon. Triplet nucleotide combinations placed immediately after the ‘TAG’ (green) that led to improved induction are underlined. Middle panel: Regulation efficiency of the constructs with various 5’UTR. Y344 with ‘CCC’ triplet nucleotide placed immediately after the ‘TAG’ led to an improved induction of 395-fold at 1 μg/ml Tc. Bottom panel: Optimizing the balance between normal splicing and drug-induced alternative splicing by randomizing the triplet nucleotide (NNN in orange) immediately after MIN5 intron. Y362 with ‘TCT’ triplet nucleotide placed immediately after MIN5 led to an improved induction of 627-fold. p<0.01, MIN5 versus Y344; p<0.01, Y344 versus Y362. **E**. The best pA regulators that combine two beneficial mutations (blue). They include an ‘A’ deletion in Arm1-1, and an ‘A’ insertion before AAUAAA. Y392, Y393, Y394, and Y362 have the ‘A’ deletion in Arm1-1 of Y359, Y360, Y361, and Y362C respectively. Y395, Y396, Y397, and Y387 have both the ‘A’ deletion in Arm1-1 and an “A” insertion immediately before AAUAAA. Statistics compare induction in fold. Data shown are average (mean ± SD) of representative experiment performed independently at least three times, each with triplicated cell cultures. Significance was determined by two-tailed Student’s t-test; n.s.: not significant, *: p<0.05, **: p<0.01, ***: p<0.001.****: p<0.0001.

To enable alternative splicing, we created and tested a series of different sequences containing the potential 3’SS ‘AG’ placed downstream of the G-quad (Fig. S9). Among these, only one construct (Y196N) gave an improved induction of 82-fold as compared to 63-fold of parental Y196. As shown in Fig. 3B, in the absence of Tc, little or no luciferase RNA was observed in both the parental Y196 (lane 1) and Y196N (lane 3) because the cleaved mRNAs are rapidly degraded. Tc treatment blocked the cleavage and led to a detectable IVS2-spliced band of 555 bp (lane 2 and 4) which was confirmed by Sanger sequencing (Fig 3B and Fig. S10A). Despite the 63-fold induction of gene expression in Y196, this band appears faint which may reflect the sub-optimal RT-PCR amplification of the stable G-quad structure. Notably, Tc treatment induced a prominent new band of 193 bp in Y196N that was absent in Y196 (lane 4 vs. 2). Sanger sequencing revealed that this band is the result of induced alternative splicing to the newly created 3’SS downstream of G-quad (highlighted as red in the sequence in Fig 3B and Fig. S10B).

To probe the effect of MAZ G-quad on alternative splicing, we altered the MAZ copy numbers from 2 to 4 in both Y196 and Y196N. This simple exchange nearly tripled the induction from 82-fold to 220-fold in Y196N. As shown in Fig. 3B, the increased induction correlates with intensified alternative splicing at the expense of IVS2 splicing (lane 4 vs. 6), indicating that 4MAZ facilitates the switching from IVS2 splicing to induced alternative splicing in Y196N. In contrast, 4MAZ did not increase but reduced the induction of Y196 from 63-fold to 45-fold, presumably because Y196 lacks the alternative splicing mechanism and 4MAZ (as opposed to 2MAZ) further enhances PAS activity making it difficult to be inhibited by Tc. Intriguingly, strengthening the alternative 3’SS by adding a branch point “TACTAAC” (S10) or a polypyrimidine tract “TCCTGTGCTT” (S11), both led to much reduced induction compared to the parental Y196N-4MAZ (Fig. 3B). Therefore, the key for this mechanism to work is a delicate balance between IVS2 and alternative splicing, so that maximal IVS2 splicing occurs in the absence of Tc and maximal alternative splicing is achieved in the presence of Tc.

To reduce the overall size of the system and the unwanted complexity of a long IVS2 intron (which may contain multiple cryptic splice sites and splicing elements), we shortened the IVS2 intron of Y196N-4MAZ from 476 to 180 nts (MIN1), and then progressively shortened it further to 120 nts (MIN4) by steps of 20 nts (detailed sequences in Fig. S11). The shortened introns showed only moderate changes in induction (Fig. 3C), indicating that much of the IVS2 intron sequence is dispensable. The construct MIN5 further mutated the two ‘ATG’ present in the MIN4 intron that may initiate unwanted protein translation, and this modification increased the induction from 233-fold to 276-fold (MIN4 vs. MIN5). Using the streamlined MIN5 as the new basis, we further fine-tuned the requirement of G-quad. As shown in Fig. 3C (right panel), five instead of four copies of MAZ did not change the induction, while six copies of MAZ reduced the induction significantly. Furthermore, replacing the MAZ with CD44 G-quad^36^ known to influence alternative splicing in the form of one, two, or four copies all reduced the induction to the minimum (Fig. 3C). These results indicate that 4MAZ is nearly optimal, and the sequence of MAZ cannot be simply substituted by other G-quad sequences.

With the intron and G-quad optimized, we then focused on other sequence properties that might affect induction. Previous reports^37, 38^ have shown that ‘unstructured’ RNA sequences placed near the start codon ATG may improve the translational efficiency, which could lead to higher protein production from each Tc-induced mRNA. To explore this property, we design a series of ‘unstructured’ RNA sequences placed before the start codon and then tested their induction. This led to Y329 with an improved 362-fold induction (Fig. 3D, unstructured sequence in blue). The results from this study also suggested that changing nucleotides immediately after the ‘TAG’ (in green, Fig. 3D), a potential alternative 3’SS in the 5’UTR, can exert a subtle influence on induction. This observation prompted us to perform a preliminary mutational analysis by randomizing the 3 nucleotides after the ‘TAG’ using an intermediate construct (see Fig. S12). The results identified a set of triplet nucleotide combinations that showed similar or improved induction (TAG, AAC, CCC, AGC, and TCT). When these triplet nucleotides were tested in Y329 as the parent construct (Fig. 3D, Y341 to Y347), the procedure led to Y344 (CCC) with an improved 395-fold induction. With the surrounding sequence of alternative 3’SS determined, we then sought to achieve an optimal balance of splicing by randomizing the triplet nucleotides immediately after the 3’SS of MIN5 intron. Fig 3D highlights four constructs (Y359 to Y362) that showed improved induction over Y344, with Y362 giving the highest induction of 627-fold at 1 μg/ml Tc.

Sequencing of Y362 also revealed an unexpected spontaneous “A” deletion in Arm1-1 (Fig. 3E and Fig. S13), which results in a ‘C’ bulge in Arm1-1. This is the same location of the “G to A” mutation that improved the performance of the earlier construct A8-103GP2 (Fig. 1F). Restoring this “A” in Y362 reduced induction (Fig. 3E, Y362C, and Fig. S13), indicating that the ‘A’ deletion yields a more favorable configuration, presumably because it destabilizes the stem near the GU-rich region. Indeed, deleting the same ‘A’ in Y359, Y360, and Y361 all consistently improved their induction (Fig. 3E and Fig. S14, Y392, Y393, and Y394 respectively). We continued to perform minor modifications including revisiting some of the earlier modifications in the 3-way junction and the arms. All of these yielded no improvement except for the “A” insertion immediately before AAUAAA, which makes the PAS more accessible in the central loop (Fig. 3E and Fig. S14, Y395, Y396, Y397, and Y387 respectively). Among them, Y395 has the highest induction of 925-fold at 1 μg/mL of Tc. The data of basal leakage expression and the induced expression used for calculating “induction in fold” of these constructs are shown in Fig. S15. Notably, Y387 shows a 735-fold induction but gives high induced expression level at 1 μg/mL of Tc, a feature that is important for applications that require high levels of induced expression (Fig. S15). This series of designs exhibit an unprecedented regulating efficiency that far exceeds the performance of previous RNA-based gene regulation systems. We have designated this “5’UTR polyA signal-based RNA switch” as the “*pA regulator*”.

## Dose response of pA regulator

The pA regulators described above were tested at 1 μg/mL of Tc (~2μM of Tc). To identify the entire regulatable range of the pA regulators, we generated dose-response curves at different drug concentrations using Y362 and Y387 as examples. As shown in Fig. 4A left panel, the dose-response curve of pA regulators plateaued at an EC_100_ (the concentration for maximum response) of 2.5 μg/ml Tc for both Y362 and Y387. At this level of maximum induction, Y362 has a regulatable range of ~900-fold and Y387 of ~800-fold. Importantly, EC_50_, the concentration for half-maximal response that defines the most sensitive part of dose-response curve, is about 0.7 μg/ml Tc for Y362 and 0.5 μg/ml Tc for Y387 (Fig. 4A, left panel). In adult humans, oral administration of Tc at the FDA-approved daily dose (500 mg twice daily) is known to yield a transient serum concentration that peaks around 5 μg/ml followed by a slow return toward a general level of 1 to 2 μg/ml^27^. Thereby, the pA regulators enable a gradient of induction with drug concentrations that fall well within the FDA-approved dose range. We also characterized the dose-response curves using the induced expression levels relative to the maximum expression level (100%) given by the parental plasmid lacking the pA regulator (HDM-Luc). As shown in Fig. 4A right panel, at EC_100_ of 2.5 μg/mL Tc, Y362 reaches about 65% of the expression level of the parental construct HDM-Luc while Y387 reaches 86%. These data indicate that Y387 can reach near full level of expression, and this feature is important for applications that require high levels of induced protein expression.

**Figure 4.**
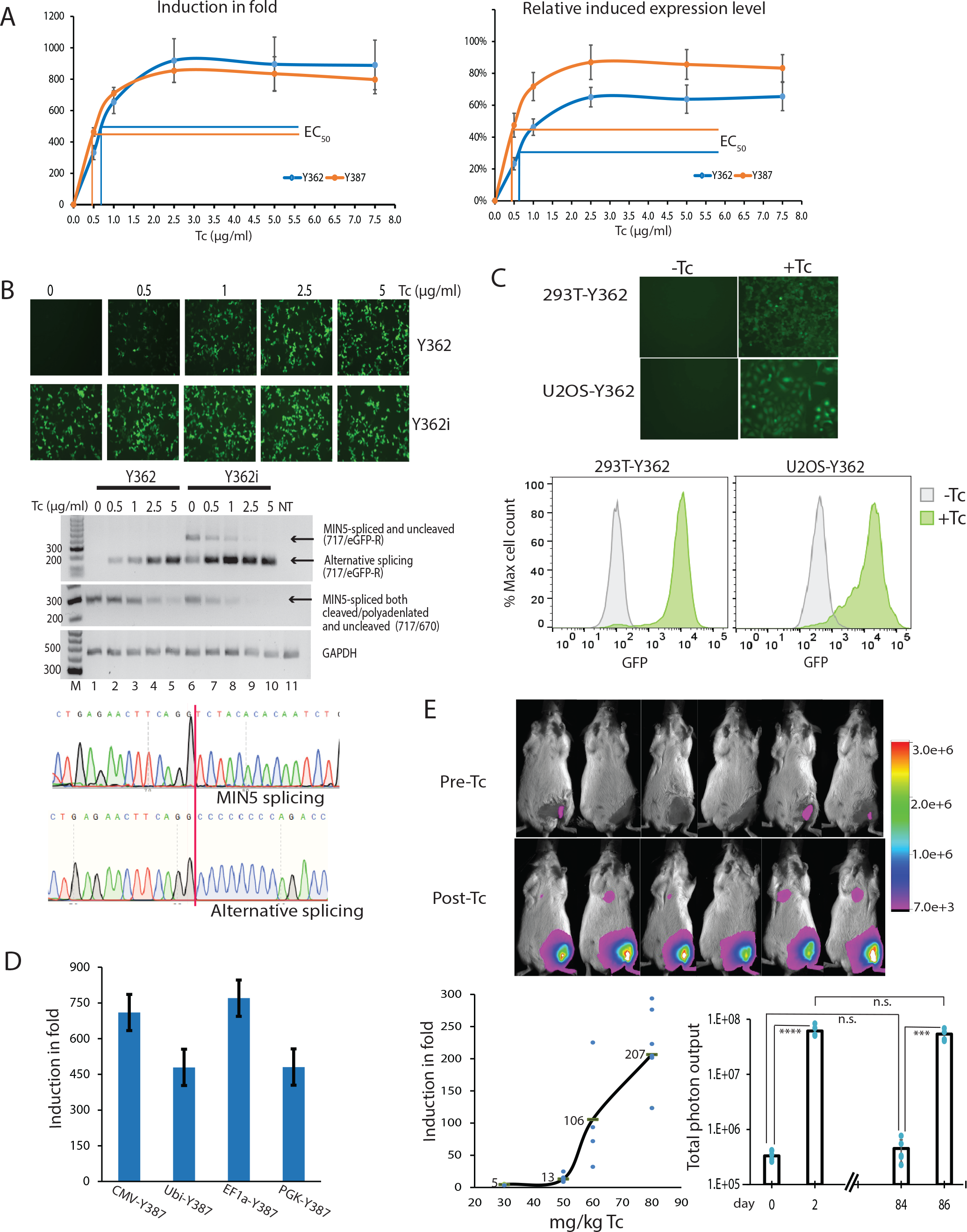
Evaluation of the pA regulator *in vitro* and *in vivo*. **A**. Dose response of Y362 and Y387 evaluated by transient transfection and luciferase assay. Left: The dose-response curve plotted based on the induction in fold. The response of pA regulators plateaued at an EC_100_ (the concentration for maximum or 100% response) of 2.5 μg/ml Tc for both Y362 and Y387 with an induction of 900- and 800-fold respectively. EC_50_, the concentration for half-maximal response, is about 0.7 μg/ml Tc for Y362 and 0.5 μg/ml Tc for Y387. Right: The dose-response curve plotted based on induced expression relative the parental construct HDM-Luc that lacks pA regulator. At EC_100_ of 2.5 μg/mL Tc, Y362 reaches about 65% of the expression level of the parental construct HDM-Luc while Y387 reaches 86%. **B**. The pA regulator works with different transgenes. 293T cells were transfected with an eGFP expressing plasmid controlled by Y362 or Y362i that has inactivated PAS, and treated with different concentrations of Tc. Top: Fluorescence microscopy images. Middle: RT-PCR analysis. Primer locations are marked by arrows in Fig. 3A. Primer pair 717/eGFP-R detects both the MIN5-spliced and alternatively spliced eGFP mRNA. Primer pair 717/670 is mainly used to detect the MIN5-spliced and polyadenylated species. GAPDH is used as loading control. With increasing Tc concentration, the expression of eGFP controlled by Y362 is increased accordingly both at the protein and the RNA levels. NT: no transfection. Bottom: Sanger sequencing confirmed the expected MIN5 splice junction in the absence of Tc, and a new alternative splice junction in the presence of Tc. The newly identified alternative 3’SS “TAG” is marked in green in Fig. 3D top panel. The red vertical line marks the splice junctions. **C**. The pA regulator functions efficiently when stably inserted into the genome of different cell lines. Top: Images of 293T and U2OS cell clones with stable Y362-eGFP insertion treated without or with 1μg/mL Tc. Bottom: Flow analysis of cell clones without (gray) or with 1μg/mL (green) Tc treatment. Histogram is plotted using the ‘Modal scale’ (FlowJo). The modal class is the bin with the highest frequency and is used as the relative 100% of cell count. **D**. The pA regulator works with different promoters. The Y387 pA regulator coupled to CMV, Ubi, EF1a, and PGK promoter were tested in 293T cells with 1μg/mL Tc and all showed efficient induction. **E**. *In vivo* evaluation of the pA regulator. Mice were injected in the gastrocnemius muscle with AAV engineered to control the luciferase reporter by the Y387 pA regulator. Top: Representative images of mice before and after Tc treatment. Background basal luciferase expression was imaged prior to Tc injection (Pre-Tc) and mice were imaged again 32 hours after Tc injection (Post-Tc) for Tc-induced luciferase expression after a single Tc injection of 80 mg/kg. Lower left: Dose-dependent response of the pA regulator *in vivo*, which was measured by ‘fold induction’ as the ratio of the Post-Tc vs. Pre-Tc luciferase signal. Lower right: The regulation by pA regulator *in vivo* is reversible. The group of mice treated with 80mg/kg Tc from day 0 to 2 was again treated with 80mg/kg Tc from day 84 to 86. Pre-Tc imaging was taken on day 0 and 84, and Post-Tc on day 2 and 86. When Tc treatment was discontinued, the luciferase expression on day 84 returned to a similar baseline level to that of day 0, and when induced again on day 86, it reaches to a similar level to that of day 2. Statistics compare the luciferase expression levels as total photon output using paired two-tailed Student’s t-test. n.s.: not significant, ***: p<0.001.****: p<0.0001. n=6 mice.

## Induced alternative splicing at the RNA level correlates with the induced protein levels

Aside from luciferase, the pA regulator functions efficiently when controlling different transgenes such as eGFP. As shown in the microscopic images in Fig 4B, 293T cells transfected with Y362 exhibited a clear dose-dependent eGFP protein expression in response to Tc. In contrast, the control construct with the PAS inactivated (AATAAA to ACACAC, Y362i) showed similarly high eGFP intensity across different Tc concentrations. To assess ligand-induced splicing, we performed RT-PCR assays using a primer pair (717/670) to detect the MIN5-spliced RNA, and another primer pair (717/eGFP-R) to detect the uncleaved therefore saved eGFP RNA species (primer locations shown in Fig. 3A). As expected, in the absence of Tc, all the detectable RNA in Y362 underwent MIN5-spliced (Fig 4B, lane 1, middle panel), and the cleavage of the synthetic PAS led to no detectable eGFP RNA in Y362 (lane 1, upper panel), mirroring what was found earlier in Fig. 3B. Increasing Tc concentrations progressively induced higher levels of alternative splicing (lane 2 to 5, upper panel, size 157 bp) at the expense of MIN5 splicing (lane 2 to 5, middle panel). This switching from MIN5 splicing to alternative splicing, which removes the synthetic PAS from RNA, correlates closely to the induced eGFP protein levels seen in the microscopic images.

In contrast, when the synthetic PAS is inactivated in Y362i without cleavage, both alternatively spliced and MIN5-spliced eGFP RNA were detected in the absence of Tc (lane 6, upper panel). This observation is consistent with the accepted notion of “co-transcriptional RNA processing”^39, 40^ that inactivated PAS weakens the splicing of last intron before it. In our configuration, it shifted the splicing balance toward the alternative splicing (lane 1 vs. 6). Again, increasing Tc concentrations progressively strengthened alternative splicing at the expense of MIN5 splicing in Y362i (lane 7 to 10, upper panel), confirming that stabilizing Y-shaped structure alone, in addition to inactivating PAS, induces alternative splicing. Sanger sequencing of the induced alternative splicing band in Y362 and Y362i revealed that the 3’SS shifted a few bases downstream from CAG to TAG in the 5’UTR (Fig 4B and Fig. S16). This shift is likely due to the new sequence added immediately after TAG (Fig. 3B), which may have strengthened TAG as the preferred alternative 3’SS.

## The pA regulator functions efficiently in different mammalian cell lines or when coupled to different mammalian promoters

In addition to transient transfection, the pA regulator also functioned efficiently when stably integrated into the genome in different mammalian cell lines. As shown in Fig. 4C, 293T and U2OS stable cell clones that carry the Y362-eGFP pA regulator showed little or no eGFP expression in the absence of Tc, and readily expressed eGFP at 1μg/mL Tc. The induced expression observed by microscopy was confirmed by flow cytometry, which showed that almost the entire cell population responded in unison to Tc (Fig. 4C). Furthermore, in addition to the default CMV promoter, the pA regulator functioned with Ubi, EF1a, and PGK promoters both in 293T cells (Fig. 4D) and HeLa cells (Fig. S17). Thus, in contrast to the Tet-On system, the pA regulator does not require any specialized promoter elements, and therefore represents a ‘portable’ motif that could be coupled to any promoter for tissue-specific applications.

## *In vivo* evaluation of pA regulator in live mice

To evaluate the potential of pA regulator in live animals, we used AAV2/9 to deliver a luciferase reporter gene controlled by the Y387 pA regulator into mice gastrocnemius muscle (the chief muscle of the hind leg) (Fig. S18). The compact size of the pA regulator allows it to be inserted in the 5’UTR of the transgene and packaged in a single viral AAV vector. An earlier *in vivo* pharmacokinetic study in mice^15^ had shown that, after a single intraperitoneal administration of 54 mg/kg Tc, a transient Tc concentration of 7.8 μM (or ~3.9 μg/ml) was detected in mouse muscle two hours after administration. Using this data as a guideline, we intraperitoneally injected mice with a single dose of 30, 50, 60 and 80mg/kg Tc to study the *in vivo* dose response. To quantify the Tc-induced luciferase expression, we first imaged the leakage baseline expression in each mouse (Pre-Tc, Fig. 4E). The induced protein expression was then imaged 32 hours after a single intraperitoneal injection of Tc (Post-Tc, Fig. 4E). The extent of induction can be calculated as the ‘fold change’ in total luciferase expression, which is the ratio of the post-Tc vs. pre-Tc luciferase signal in the same animal. As shown in Fig. 4E, before the Tc treatment, mice showed very little leakage baseline expression. In contrast, strong luciferase signals were induced after a dose of 80mg/kg Tc. Quantitative analyses based on total photon output showed that gene expression was induced on average to 5, 13, 106 and 207-fold when mice were injected with 30, 50, 60 and 80mg/kg Tc respectively (Fig. 4E, lower left), indicating that the induction *in vivo* also followed a dose-dependent pattern similar to that of cultured cells. Under the highest single dose we tested (80mg/kg Tc), the pA regulator can effectively turn on luciferase expression up to 300-fold in some mice. Moreover, the induced gene expression by pA regulator is reversible *in vivo* (in terms of both the baseline expression and the induced levels). This was demonstrated by redosing the same group of mice separated by a prolonged period. As shown in Fig. 4E lower right, the pre-Tc baseline expression on day 84 returned to a similar level to that of pre-Tc on day 0, while the induced luciferase expression measured on day 86 reached a level comparable to that measured on day 2. Together, the results demonstrate that gene expression under the control of the pA regulator in live animals is both dose-dependent and reversible, and that repeated induction can be achieved with long-term stability *in vivo*.

## Discussion

The engineering principles used to construct the pA regulator symbolize a radical departure from the conventional riboswitch approaches. First, while natural riboswitch utilizing a PAS in the 3’ UTR was found in *Arabidopsis thaliana*^41^ and PAS has been suggested as a gene control tool^42, 43^, the pA regulator we described positions the PAS in the 5’UTR, a unnatural location for mammalian transcripts. Yet the cleavage efficiency limits the basal leakage expression to 0.1%, surpassing that of self-cleaving N79 ribozyme developed by the author previously^12^. Second, numerous systems using a single cb32 aptamer achieved only limited induction^4, 17^, indicating that ligand binding to a single aptamer is insufficient to produce the amount of energy change required for efficient switching. We overcame this limitation by inventing the Y-shaped structure that effectively combines the effects from multiple aptamers, in addition to the use of ‘A42G’ mutant aptamer that improves switching efficiency. Third, most published RNA-based gene regulation systems have employed only a single control mechanism. In contrast, the pA regulator is governed by two distinct and complementary mechanisms: (a) direct clamping of the PAS by Tc binding, and (b) induced alternative splicing that permanently removes the PAS. In fact, alternative splicing removes nearly all the elements that constitute the pA regulator (the intron, Y, PAS, and G-quad) from each RNA, resulting in a mature mRNA with a 5’UTR sequence closely mirroring the parent gene 5’ UTR. The strategy of ‘dual mechanism’ also ensures maximal control and minimizes potential performance variability in different cell types due to differences in cellular context. Lastly, the Y-shaped structure of pA regulator is engineered with strong RNA secondary structures. Ligand binding to the aptamers involves stabilizing the Y-shaped structure and no complicated RNA folding strategies such as helix slippage^44^ or the use of communication module sequences^45^ are involved in the mode of action. As such, the pA regulator represents a modular platform that could allow simple plug-and-play exchange of the Tc aptamer by alternative and diverse aptamers^46^ with minimum modifications. This represents important features because *in vivo* gene regulation applications are often hindered by limited tissue penetrance of the inducer ligands. The flexibility to accommodate new aptamers specific to ligands with desirable pharmacokinetic properties would significantly broaden future applications targeting different tissues. These engineering principles, each indispensable to the overall efficiency and practicality, are integrated as a single functional pA regulator unit that achieves an unprecedented level of regulation.

Unlike the Tet-On system that requires one promoter to generate the regulatory transactivator protein and a second promoter to express the transgene, the pA regulator is a compact single promoter system that can be implemented in a single plasmid or a single viral vector. In contrast to the ‘irreversible’ and “On or Off two-stage only” gene control generated by CRISPR/Cas9 DNA editing, the pA regulator controls gene expression in a “dose-dependent’ and “reversible” manner, both features crucial for effective therapeutics^3^ as well as the biological study of gene function. In addition, the CRISPR/Cas9 system is known to generate high immunogenicity in hosts^47, 48^. Compared to systems based on alternative splicing such as the X-on switch^20^ that generate additional amino acid sequences therefore promoting transgene-specific immune response, the pA regulator requires no change to the transgene coding sequence.

In summary, gene regulation in clinical settings requires demanding features of non-immunogenicity, high dynamic ranges, low leakage expression, and low ligand concentrations for induction. The pA regulator described here represents an important milestone in mammalian gene regulation technology to fulfill such needs and could open a new window of opportunity in therapeutics as well as biological studies.

## Acknowledgements

We thank Tom Cooper and Richard Sifers for critical suggestions. We thank the Gene Vector Core at Baylor College of Medicine and Dr. Kazuhiro Oka for consultation and AAV production.

## Authors contributions

L.L., J. D.-Y. J, Y.W., and P.-W. C performed experiments. L.Y. conceived the project and obtained the fundings. L.L., J. D.-Y. J, and L.Y. wrote the manuscript.

