## Supplementary materials for "pA regulator: a system for controlling mammalian gene expression via the modulation of polyA signal cleavage"

**Summary**:

The pA regulator is the first clinically relevant gene regulation system with a ligand $EC_{50}$ concentration approved by FDA.

**Supplementary Materials:**

### pA regulator: a system for controlling mammalian gene expression via the modulation of polyA signal cleavage

Liming Luo, Jocelyn Duen-Ya Jea, Yan Wang, Pei-Wen Chao, Laising Yen

Department of Pathology & Immunology, Department of Molecular and Cellular Biology, Dan L. Duncan Cancer Center, Baylor College of Medicine, Houston, TX 77030, USA

#### Materials and Methods

##### Luciferase assay in mammalian cells

Cells were seeded in 96-well plates at a density of 2.5 to $3 \times 10^4$ cells/well. After 24 hours of incubation, each well was transfected with 50 ng of DNA vectors and was incubated with culture medium (DMEM, Gibco, Cat#11995073) with 10% fetal bovine serum (Corning, Cat#35010CV) and 10% Penicillin-Streptomycin Solution (Corning, Cat#30001CI), without or with various concentration of tetracycline for an additional 18 hours. Luciferase activity was measured in relative light units (RLU) with the Polarstar Omega plate reader (BMG Labtech). To make 36 mL of assay buffer, 144 µL 1M DTT, 108 µL 0.1 M ATP, 252 µL 0.1M luciferin and 360 µl 0.05M CoA were added to 35 mL of basic buffer (25mM Tricine, 0.5mM EDTA-Na2, 0.54mM Na-triphosphate, 16.3 mM MgSO4.7H2O, and 0.8% Triton X-100). After the cell medium was removed, 40 µL of assay buffer was added to each well, and luciferase activity was read twice with the Polarstar Omega plate reader.

##### RT-PCR

Cells transfected with the respective constructs were cultured for 18 hours at 37 °C in medium in the absence or presence of tetracycline. Total RNA was isolated according to the protocol supplied with RiboPure™ RNA Purification Kit (Ambion, Cat#AM1924). For RT-PCR, RT was performed using SuperScript III (Invitrogen, Cat#18080044) according to the manufacturer's protocol and PCR was performed using the KOD DNA polymerase (Milliporesigma, Cat#71085). PCR primers used for amplifying pA regulator RNA are: 717: 5'- TCAGATCGCCTGGAGACGCC-3', 670: 5'- TTTGAATTCGATCTGGTATGTTTTGCCACAAACAC-3', Luc-R: 5'-_TCTTCCAGCGGATAGAATGG-3', eGFP-R: 5'- GTGAACAGCTCCTCGCCCTT-3'. PCR primers used for amplifying the GAPDH RNA include GAPDH F1: 5'-GCGTCTTCACCACCATGGAGA -3', GAPDH R1: 5'- AGCCTTGGCAGCGCCAGTAGA -3'.

##### Stable cell line generation

Cells were seeded in 6-well plates at a density of $2.4 \times 10^5$ cells/well. Twenty-four hours after seeding, each well was transfected with a mixture of the respective plasmid and puromycin plasmid at a ratio of 5:1. Cells were incubated for an additional 48 hours, and then cultured with fresh medium containing 1µg/mL puromycin to select single clones with stable genomic insertion. Puromycin resistant clones were selected and checked for the expression of the desired plasmid.

##### Fluorescence Microscopy

For simple transfection, cells were seeded in 12-well plates at a density of $1.2\times 10^5$ cells/well. After 24 hours of incubation, each well was transfected with 500 ng of DNA vectors and were incubated with culture medium containing none or various concentrations of tetracycline for an additional 18 hours. For stable cell lines, cells were incubated with culture medium containing none or various concentration of tetracycline for 48 hours. Images were taken on a fluorescence microscope (Zeiss Axiovert 40CFL) at a magnification of 200x.

**Flow analysis**

Puromycin-selected pA regulator single clones were seeded in 12-well plates at a density of $1.2\times 10^5$ cells/well. After 24 hours of incubation, each well was incubated in the presence (1 µg/ml) or absence of tetracycline for 24 hours and then harvested for flow analyses. pA regulator-controlled GFP expression levels of single clones were quantified using LSR Fortessa (BD biosciences) and analyzed by FlowJo software (FlowJo).

**Generation of adeno-associated virus vector**

Y387 pA regulator was cloned into the 5′ UTR in pAAV.CMV.ffLuciferase.SV40 plasmid between SacI and ApaI restriction sites. Subsequently, the luc2 of the original vector was replaced by firefly luciferase sequence between NotI and ClaI sites (see Fig. S18 for AAV diagram). All restriction enzymes are from New England Biolabs. AAV-Y387-luc (AAV2/9) virus (titer: 1.84 x $10^{14}$ genome copy/ml) was produced by the Gene Vector Core at Baylor College of Medicine. pAAV.CMV.ffLuciferase.SV40 was a gift from James M. Wilson (Addgene plasmid #105532; http://n2t.net/addgene:105532 ; RRID:Addgene_105532).

**pA regulator-controlled luciferase expression in mice**

Stock pAAV-Y387-luc AAV was diluted to 1.5 x $10^9$ /µl for intramuscular injection. 40 µl AAV solution (6 x $10^{10}$ AAV particle) was injected into medial gastrocnemius muscle in the right hind leg of 8 six-week-old female ICR mice (Fig. S18). Tetracycline Hydrochloride (Sigma, Cat#T7660) was dissolved in physiological saline (VWR, Cat#470302-026) to 3, 5, 6, or 8 mg/ml and injected into mice at 10 µl/g bodyweight through intraperitoneal (IP) injection. Luciferase expression levels in the mice were measured before (Pre-Tc) and at 32 hours after Tetracycline treatment (Post-Tc). The induction in fold was calculated by the ratio of Tc-induced luciferase expression to baseline expression. Experimental timeline is shown in Fig. S18. The dosage response of pA regulator *in vivo* was determined by induction of the same mice under single dose treatment at 30, 50, 60, and 80 mg/kg Tc respectively. Each treatment is separated by 3 to 4 weeks to allow luciferase degradation and return to baseline background signals.

***In vivo* bioluminescent imaging**

To measure the luciferase expression, mice were injected with luciferase substrate (D-Luciferin, GoldBio, Cat#LUCNA) by IP injection. D-Luciferin was dissolved in DPBS (Corning, Cat#21-040-CV) at 15 mg/ml, filtered through 0.2 μm filter, and injected into mice by 10 μl/g bodyweight. After 5 minutes of luciferin injection, baseline and Tc-induced luciferase expression in mice were imaged by Bruker XP Pro system (Bruker) with 10 minutes of exposure time. Due to the limitation of the imaging system, luminescence signals over 160-fold differences cannot be processed using the same exposure setting, because the Tc-induced expression would exceed linear range of the software. To acquire accurate luciferase readings, we captured pre- and post-Tc treatment mice images by different binnings and normalized their signals to the average radiance (photons/sec/$cm^2$). Baseline luciferase expression was imaged by 4x4 bins whereas induced expression was imaged with 1x1 bins to avoid photon saturation. Normalized luminescence signals were measured by Fiji (ImageJ) to calculate Tc-induced fold induction in each mouse.

**Supplementary Figures**

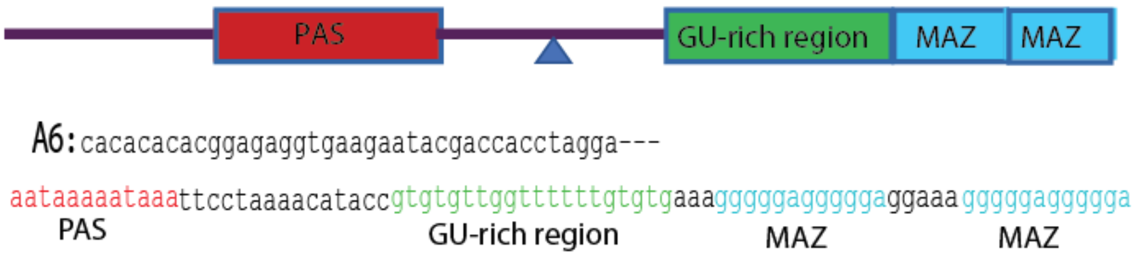


**Fig. S1. Downstream elements used to enhance the function of synthetic polyA signal (PAS) at the 5'UTR.** A modified GU-rich region and 2 copies of G-rich motif called MAZ (that constitutes a typical G-quad sequence) are placed downstream of PAS to enhance its cleavage. The construct A6 is used as an example to show the sequence of downstream elements. Red: PAS. Green: GU-rich region. Blue: MAZ motif. Triangle: Cleavage site that lies between the PAS and the GU-rich region.

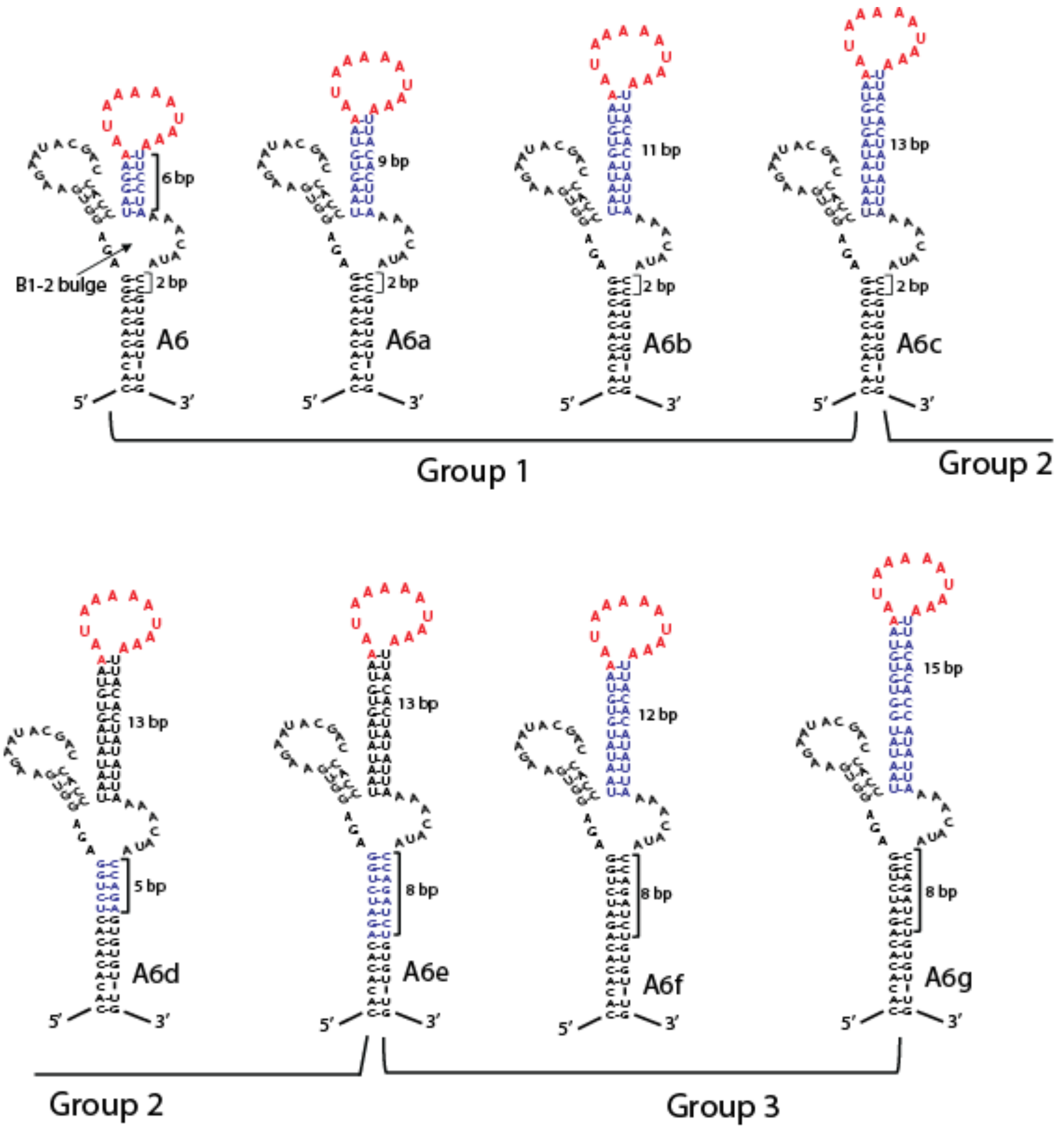


**Fig. S2. Sequences of A6a to A6g used to probe the effects of P1 stem length and the position of GU-rich region on induction.** Group 1 (A6 to A6c): P1 stem length varied from 6 to 13bp while the GU-rich region distance to the B1-2 bulge fixed at 2bp. Group 2 (A6c to A6e): the distance of GU-rich region to the B1-2 bulge varied from 2 to 8 bp while the P1 stem length fixed at 13bp. Group 3 (A6e to A6g): P1 length varied from 12 to 15 bp while the GU-rich region distance to the B1-2 bulge fixed at 8bp. Red: PAS. Blue: areas of modification.

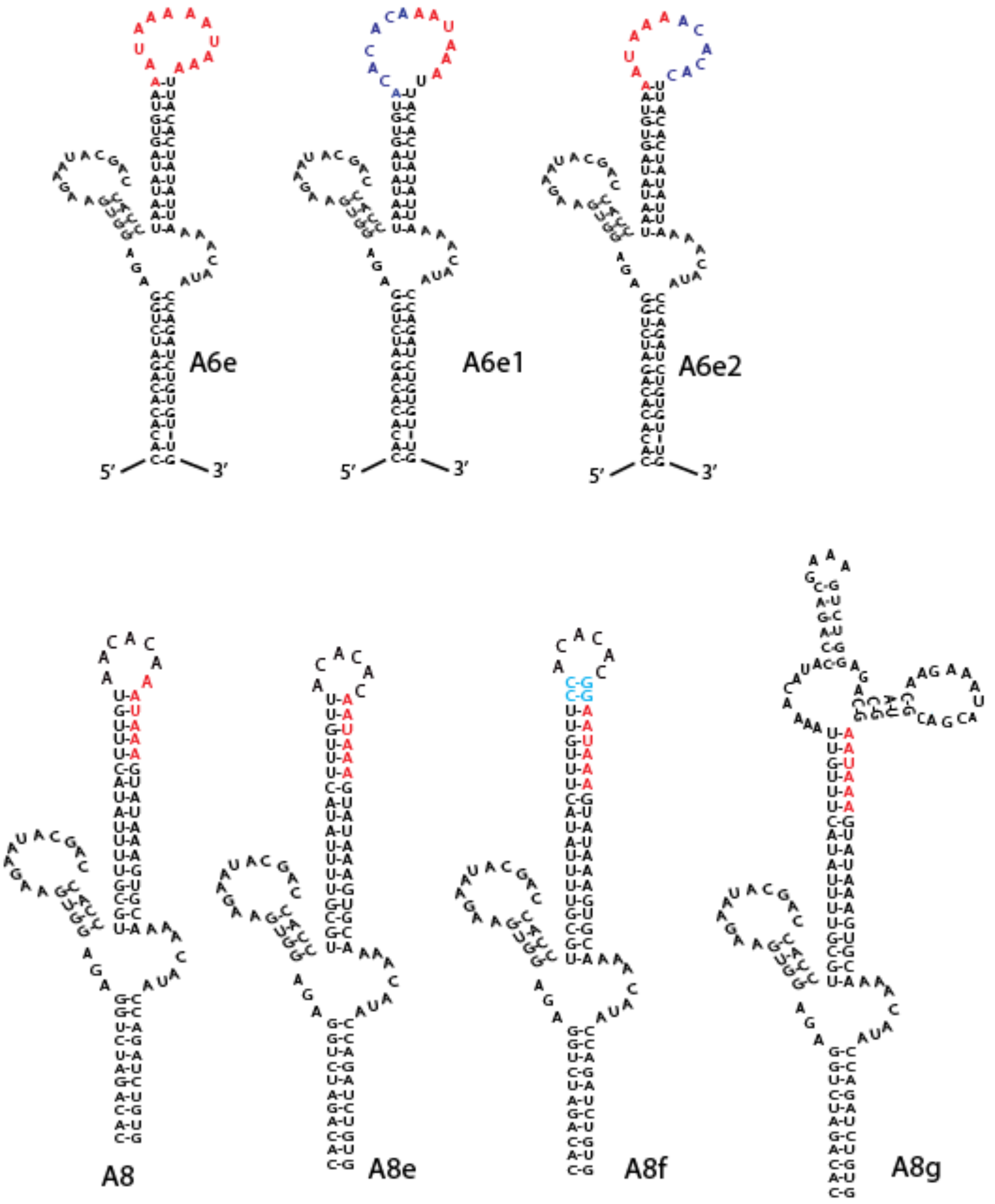


**Fig. S3. Sequences of A6e to A6e2 used to determine individual PAS contribution to the overall cleavage efficiency, and sequences of A8 to A8g used to determine the effects of PAS clamping.** A6e1: the first PAS is mutated. A6e2: the second PAS is mutated. Blue: mutated PAS. A8: PAS was partially embedded in P1 stem; A8e: PAS was completely embedded in P1 stem; A8f: PAS was completely embedded in P1 and clamped with 2 additional C-G pairs (light blue); A8g: PAS was completely embedded in P1 and clamped by an additional Tc aptamer. Red: PAS.

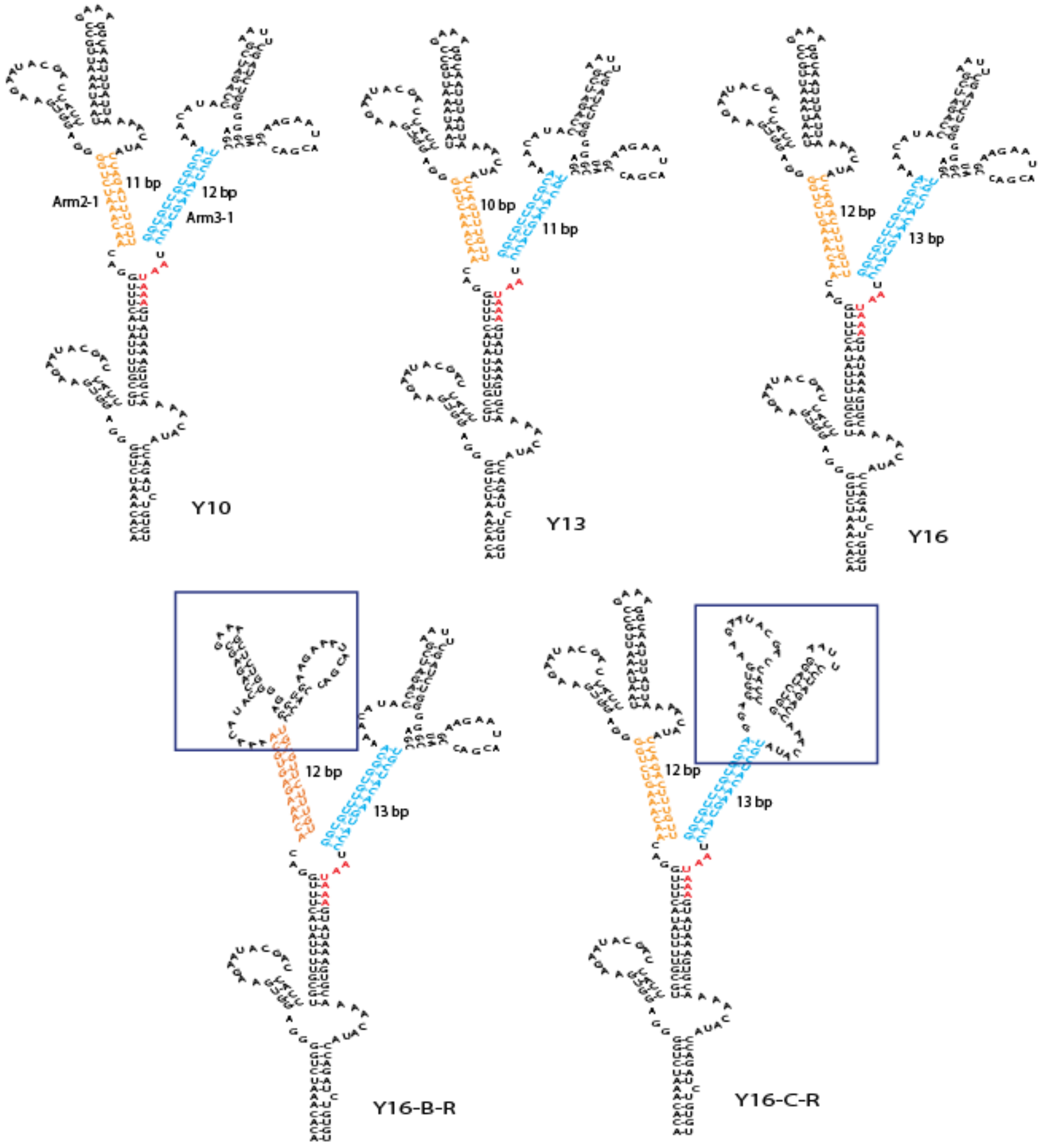


**Fig. S4. Sequence of constructs with the Y-shape structure.** Y10, Y13, and Y16 have Arm2-1 or Arm3-1 ranging from 10 to 13 bp. Y16-B-R has aptamer B in the reverse orientation as opposed to the forward orientation of Y16. Y16-C-R has aptamer C in the forward orientation as opposed to the reverse orientation of Y16. Red: PAS. Orange: areas of modification in Arm2-1. Blue: areas of modification in Arm3-1. Box: the inverted aptamer.

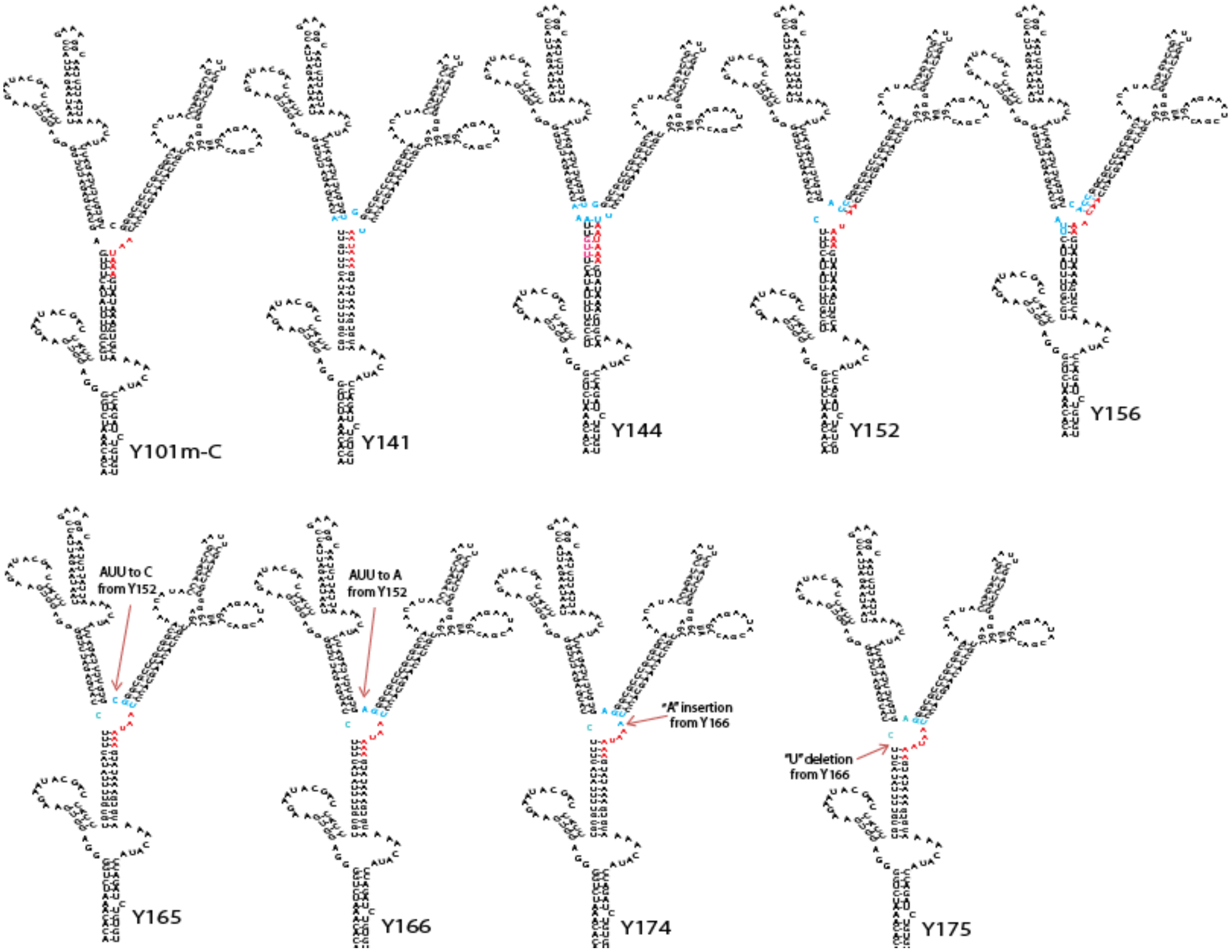


**Fig. S5. Sequence of the constructs used to optimize the location of PAS within the 3-way junction.** Y141 and Y144: PAS was moved further into Arm1-2 relative to that of Y101m-C. Y152 and Y156: PAS was moved further into Arm3-1 relative to that of Y101m-C. Y165 and Y166: "AUU" was replaced by a single base (arrow) from Y152. Y174: an "A" was inserted (arrow) before AAUAAA from Y166. Y175: a "U" was deleted (arrow) from Y166. Among this group, Y175 struck an optimal balance with 4 nts of PAS in the central loop and 2 nts embedded in Arm1-2. Red: PAS. Blue: areas of modification.

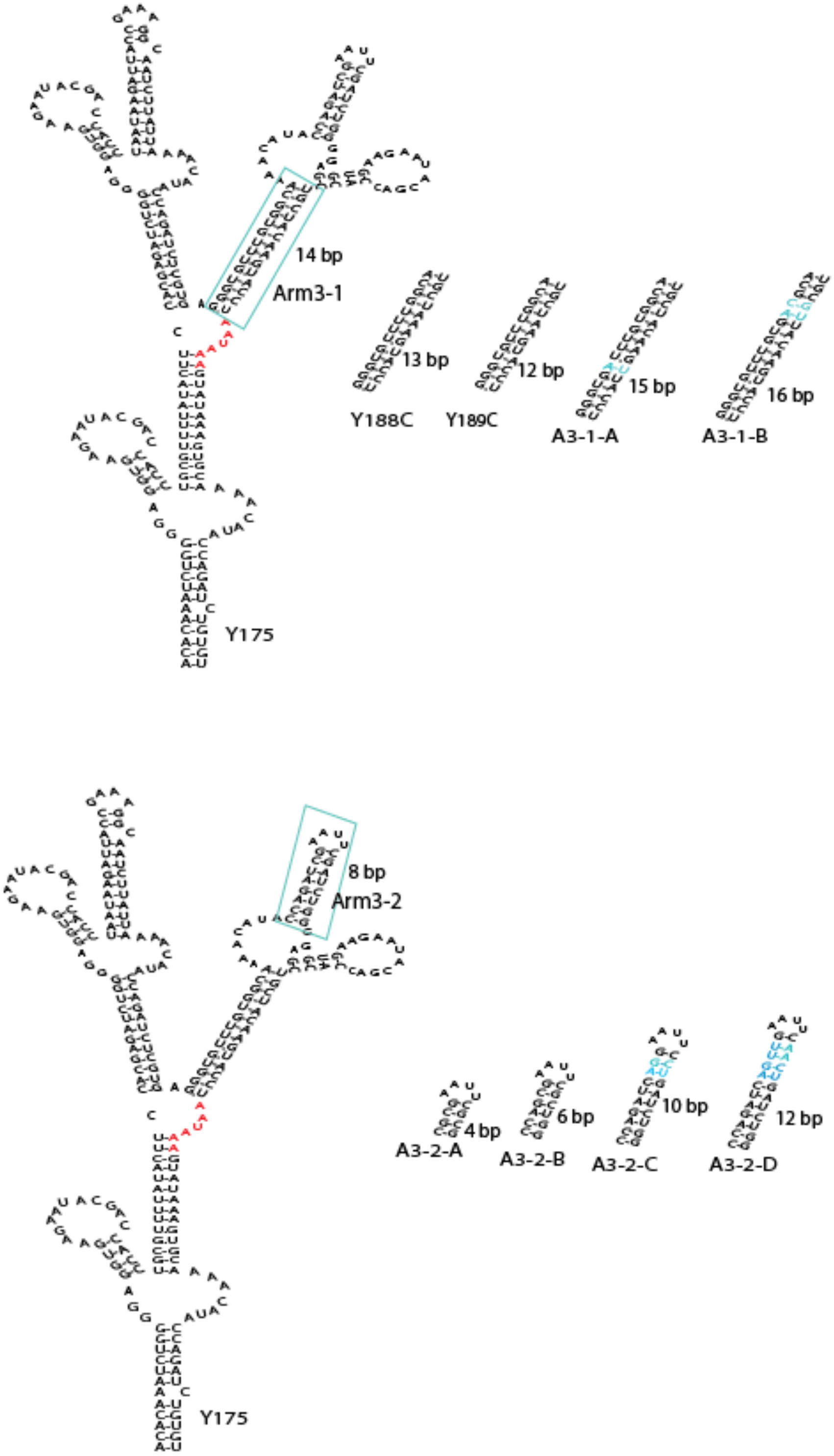


**Fig. S6. Sequence of the constructs derived from Y175 with Arm3-1 and Arm3-2 modifications.** Upper: The length of Arm3-1 was shortened to 13 bp (Y188C) and 12 bp (Y189C), or increased to 15 bp (A3-1-A) and 16 bp (A3-1-B). Lower: The length of Arm3-2 was shortened to 4 bp (A3-2-A) and 6 bp (A3-2-B), or increased to 10 bp (A3-2-C) and 12 bp (A3-2-D). Red: PAS. Blue: areas of modification.

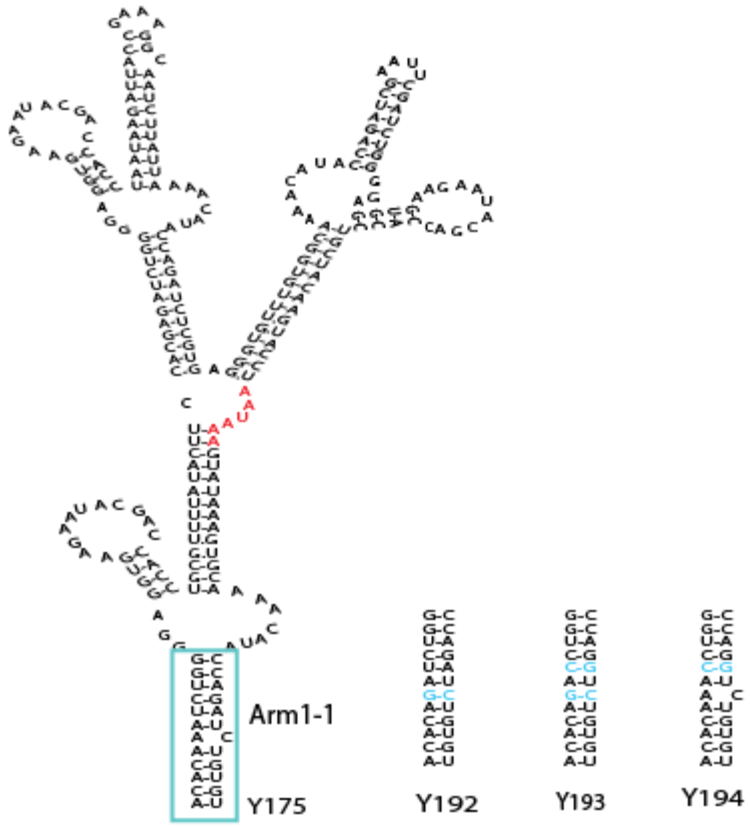


**Fig. S7. Sequence of the constructs derived from Y175 with Arm1-1 modifications.** Y192 has 'A' to 'G' change to close the unpaired break in Arm1-1, making arm1-1 more stable. Y193 has the same change as Y192 and a "U-A" pair to "C-G" in Arm1-1, so Y193 is more stable than Y192. Y194 has "U-A" to "C-G" change only in Arm1-1 but no 'A' to 'G' change to close the unpaired break. Red: PAS. Blue: areas of modification.

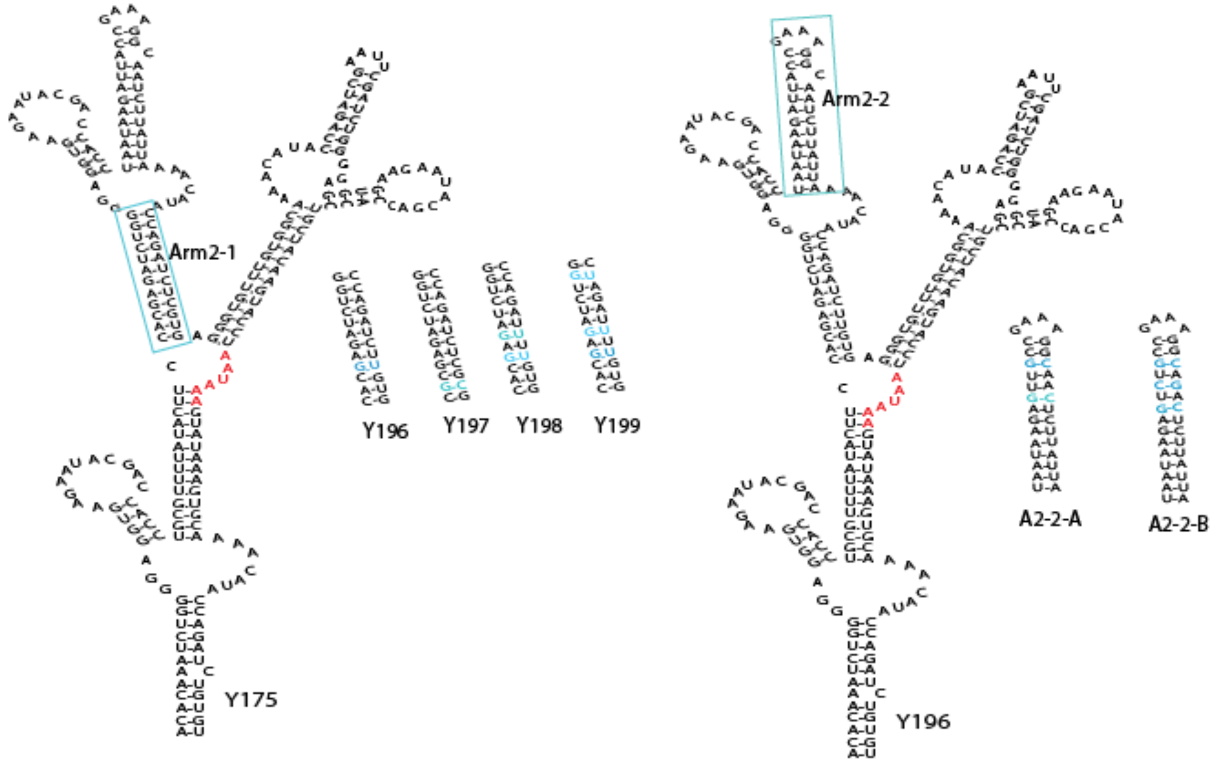


**Fig. S8. Sequence of the constructs derived from Y175 with Arm2-1 modifications, and derived from Y196 with Arm2-2 modifications.** Y196 has a "G-C" to "G-U" change making Arm2-1 less stable compared to Y175. Y197 has an "A-U" to "G-C" change making Arm2-1 more stable compared to Y175. Y198 and Y199 both have additional "G-C" to "G-U" changes as compared to Y196, making Arm2-1 even less stable. A2-2-A has 'A' to 'G' change to close the unpaired break in Arm2-2, and a "G-C" pair insertion to make Arm2-2 more stable. A2-2-B has one additional "C-G" pair insertion to make Arm2-2 more stable. Red: PAS. Blue: areas of modification.

| Construct | 5'UTR | Induction in fold (1 μg/mL Tc) |
|---|---|---|
| S72 | GCGGCCGCCaacaacaacaacaattcctgctcctcttctgcccaggaacacgcttgccttccccaaggcttccagaagctctgaggcaggaggcaccaagttctacctcacgtttggaggatcttgctagctATG | 3 |
| S73 | GCGGCCGCCagcagatccagtgcttcctgctcctcttctgcccaggaacacgcttgccttccccaaggcttccagaagctctgaggcaggaggcaccaagttctacctcacgtttggaggatcttgctagctATG | 5 |
| S76 | GCGGCCGCCaacaacaacaacaattcctgctcctcttctgccctggaacacgcttgccttccccaaggcttccagaagctctgaggcaggaggcaccaagttctacctcacgtttggaggatcttgctagctATG | 16 |
| S77 | GCGGCCGCCagcagatccagtgcttcctgctcctcttctgccctggaacacgcttgccttccccaaggcttccagaagctctgaggcaggaggcaccaagttctacctcacgtttggaggatcttgctagctATG | 8 |
| S83 | GCGGCCGCCaacaacaacaacaacaacaacaacaacacgcttgccttccccaagcttccacaagcaacctcaaacagacaccATG | 13 |
| S84 | GCGGCCGCtactaacaacacgcttgccttccccaagcttccacaagcaacctcaaacagacaccATG | 4 |
| S85 | GCGGCCGCgcttcctgctcctcttctgcccaggaacacgcttgccttccccaagcttccacaagcaacctcaaacagacaccATG | 6 |
| S86 | GCGGCCGCgcttcctgctcctcttctgccctggaacacgcttgccttccccaagcttccacaagcaacctcaaacagacaccATG | 8 |
| S87 | GCGGCCGCTactaacgctcctcttctgccctggaacacgcttgccttccccaagcttccacaagcaacctcaaacagacaccATG | 12 |
| S89 | GCGGCCGCtactaacgctcctcttctgcccaggaacacgcttgccttccccaagcttccagaagcaacctcaaacagacaccATG | 3 |
| S90 | GCGGCCGCCaacaacaacaacaacaacaacaacaacaacaacaacaactactaacataacagtgttcactagcaacctcaaacagacaccATG | 22 |
| S91 | GCGGCCGCCaacaacaacaacaacaacaacaacaacaacaacaacaactcctgtgcttataacagtgttcactagcaacctcaaacagacaccATG | 23 |
| Y196N | GCGGCCGCCcaacaacataacagtgttcactagcaacctcaaacagacaccATG | 82 |

**Fig. S9. Designed and tested 5’UTR sequences containing a potential alternative 3’ splice site ‘AG’ (red) placed downstream of the G-quad.**

A

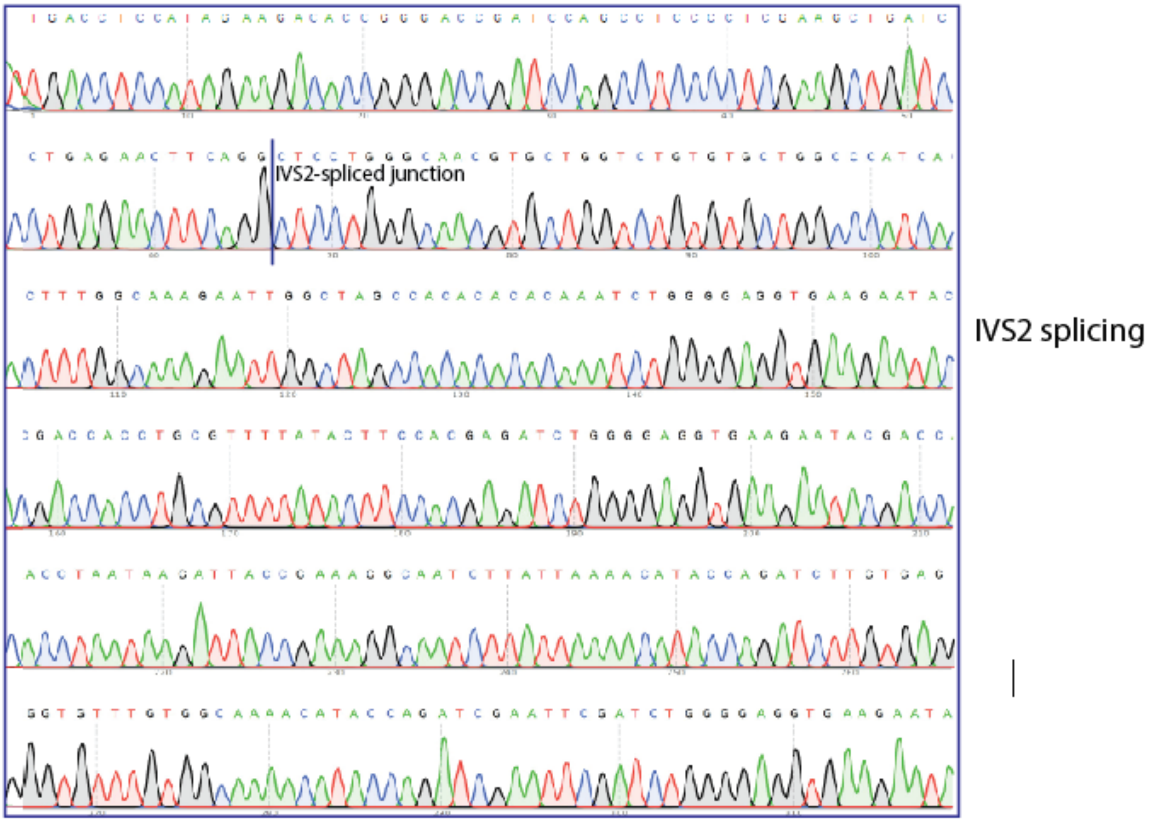


Intron sequence (in red ) near exon junction: TTCAGG[GTgagt……cccacAG]CTAGCC

B

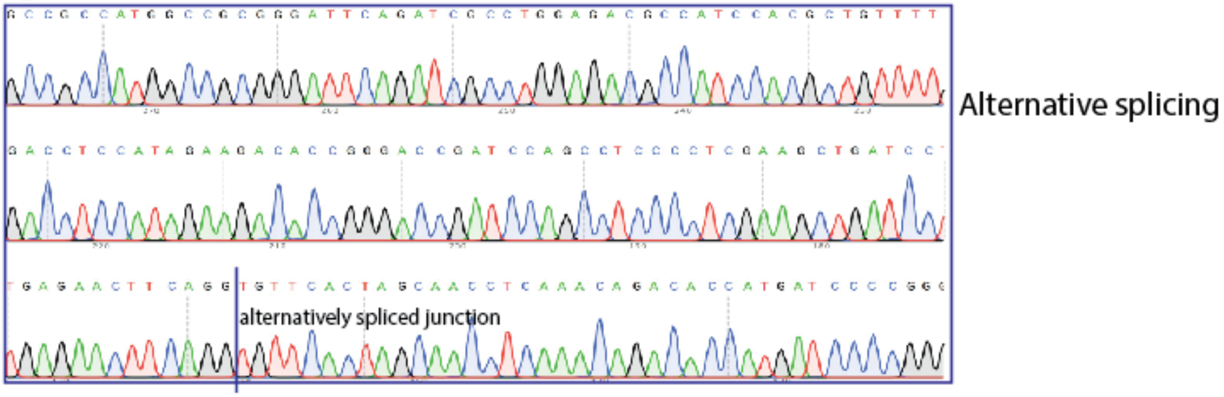


Intron sequence (in red) near exon junction: TTCAGG[GTgagt ……ataacAG]TGTTCAC

**Fig. S10. Sanger sequencing confirmation of splicing in Y196N-4MAZ.** A: Sanger sequencing of IVS2-spliced band. B: Sanger sequencing of the alternatively spliced band. The splice junction is indicated by a vertical line. The intron sequence near the exon junction is in red.

```
MIN1: Intron Size 180 bases
GTgagtctatgtttctgcatataaattgtaactgatgtaagaggtttcatattgctaatagc
agctacaatccagctaccattctgcttttattttatggttgggataaggctggattattctg
agtccaagctaggcccttttgctaatcatCttcatacctcttatcttcctctgCAG

MIN2: Intron Size 160 bases
GTgagtctatgctgatgtaagaggtttcatattgctaatagcagctacaatccagctaccat
tctgcttttattttatggttgggataaggctggattattctgagtccaagctaggcccttttt
gctaatcatcttcatacctcttatcttcctctgCAG

MIN3: Intron Size 140 bases
GTgagtctatgttgctaatagcagctacaatccagctaccattctgcttttattttatggtt
gggataaggctggattattctgagtccaagctaggcccttttgctaatcatcttcatacctc
ttatcttcctctgCAG

MIN4: Intron Size 120 bases
GTgagtctatgccagctaccattctgcttttattttatggttgggataaggctggattattc
tgagtccaagctaggcccttttgctaatcatcttcatacctcttatcttcctctgCAG

MIN5: Intron size 121 bases
GTgagtcttaagccagctaccattctgcttttattttatcgttgggataaggctggattatt
ctgagtccaagctaggcccttttgctaatcatcttcatacctcttatcttcctctgCAG
```

**Fig. S11. The sequences of MIN1 to MIN5 introns derived from IVS2 intron.** IVS2 intron was shortened from 476 nt to 180 nt (MIN1) first, and then the size was reduced by 20 nt stepwise from MIN2 to MIN4. MIN5 (121 nt) has the same sequence as MINI4 but with two unwanted AUG mutated (in blue).

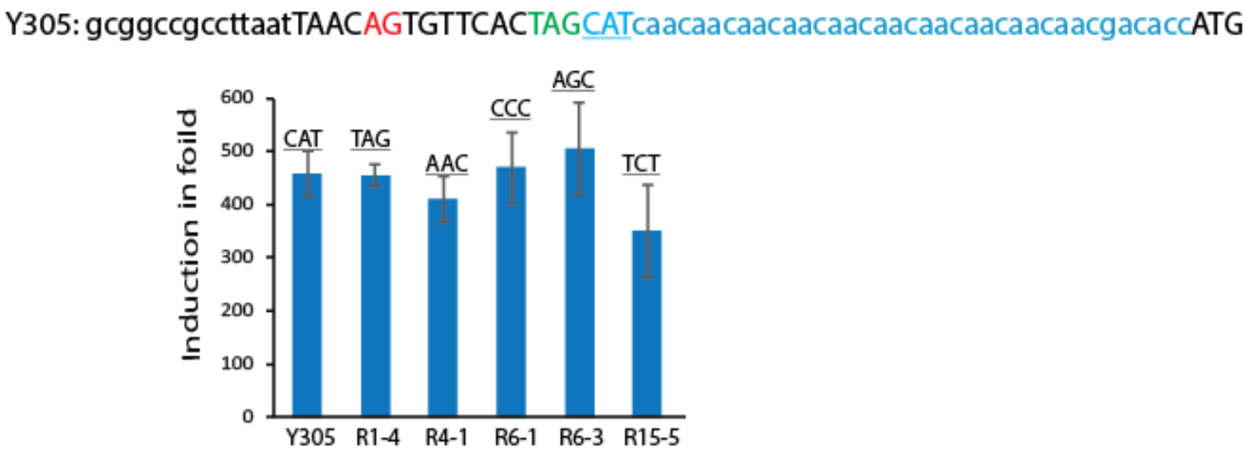


**Fig. S12. A set of triplet nucleotide combinations that showed similar or improved induction (TAG, AAC, CCC, AGC, and TCT) were identified by randomizing the 3 nucleotides after the 'TAG' in Y305.** Top: The 5'UTR before the start codon in Y305. The randomization location is underlined.  Bottom: Regulation efficiency of the constructs with various triplet nucleotide combinations.

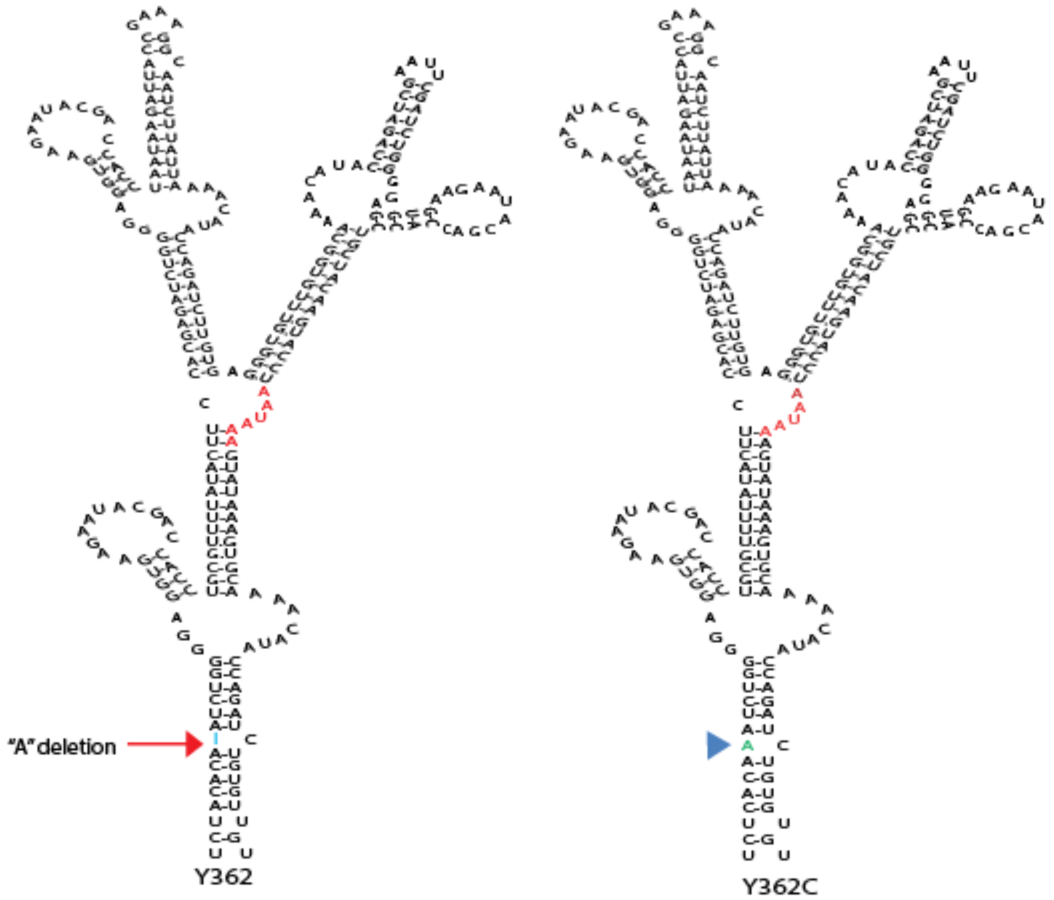


**Fig. S13. A spontaneous 'A' deletion was identified in Arm1-1 of Y362.** When this "A" was inserted back the original location (blue arrow in Y362C), the induction was significantly decreased.

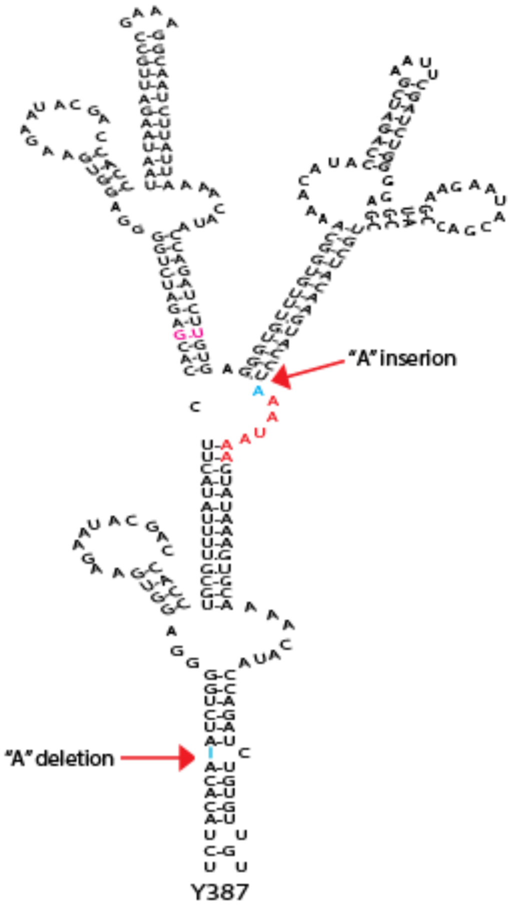


| | Y359 | Y392 | Y395 | Y360 | Y393 | Y396 | Y361 | Y394 | Y397 | Y362C | Y362 | Y387 |
|---|---|---|---|---|---|---|---|---|---|---|---|---|
| "A" deletion in Arm1-1 | No | Yes | Yes | No | Yes | Yes | No | Yes | Yes | No | Yes | Yes |
| "A" insertion before AAUAAA | No | No | Yes | No | No | Yes | No | No | Yes | No | No | Yes |
| Bases after MIN5 | CAU | CAU | CAU | UUU | UUU | UUU | UGA | UGA | UGA | UCU | UCU | UCU |
| Induction in fold | 472 | 631 | 925 | 407 | 555 | 775 | 424 | 483 | 922 | 260 | 627 | 735 |

**Fig. S14. The combined effect of single base deletion in Arm1-1 and single base insertion in 3-way junction.** Top: Y387 as an example showing single base changes at the two indicated locations. Bottom: 12 constructs designed to probe the cumulative effects of these changes.

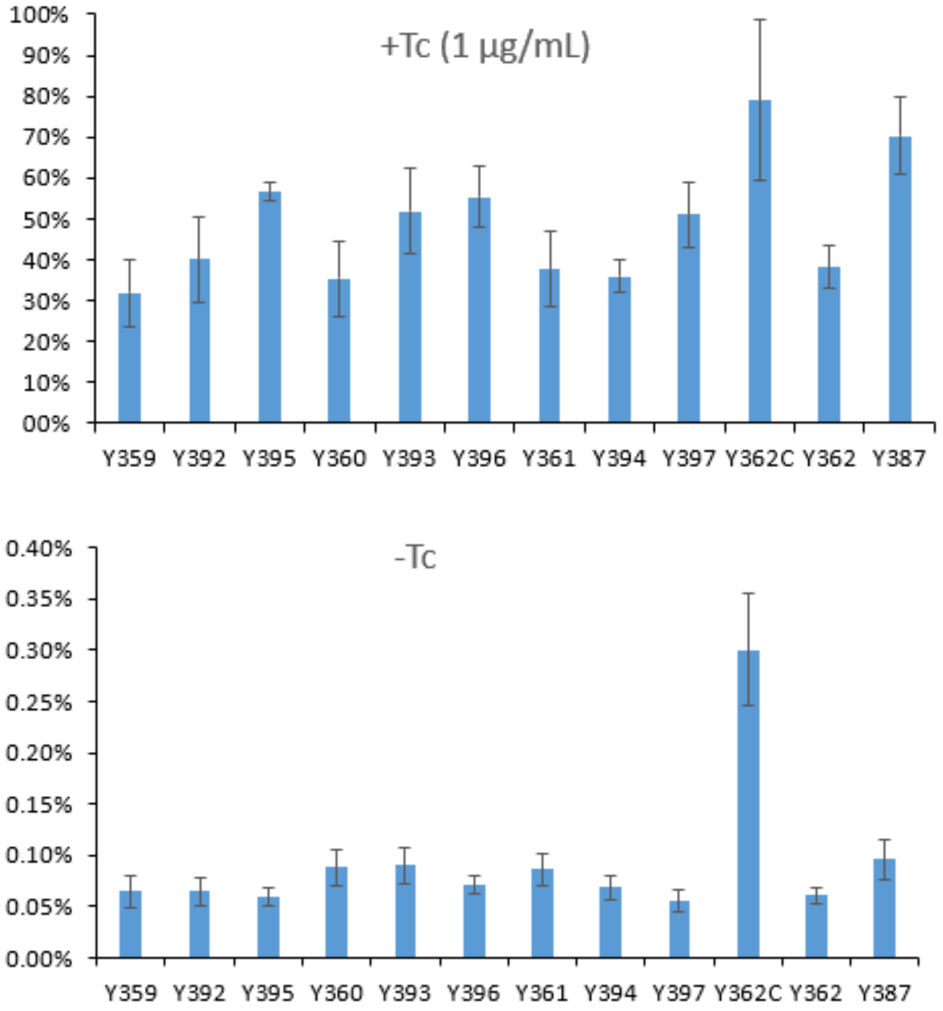


**Fig. S15.** Induced expression levels (+Tc) and basal leakage background expression (-Tc) of the pA regulator constructs listed in Fig. S13. The expression levels are normalized to that of the parental construct HDM-Luc lacking the pA regulator. Top: Tc-induced expression level. Bottom: Leakage background expression without Tc treatment.

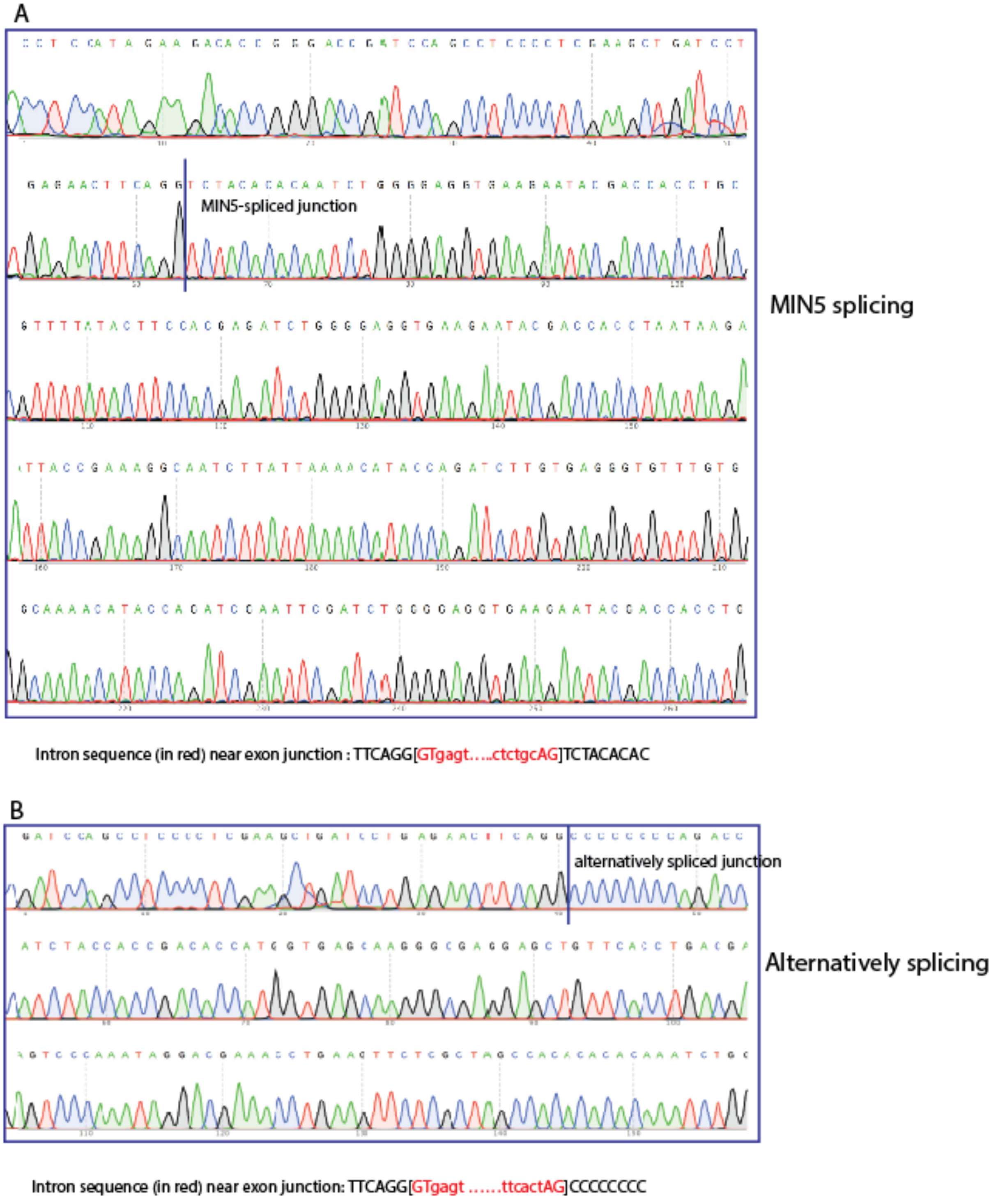


**Fig. S16. Sanger sequencing confirmation of splicing in Y362-eGFP.** A: Sanger sequencing of MIN5-spliced band. B: Sanger sequencing of the alternatively spliced band. The splice junction is indicated by a vertical line. The intron sequence near the exon junction is in red.

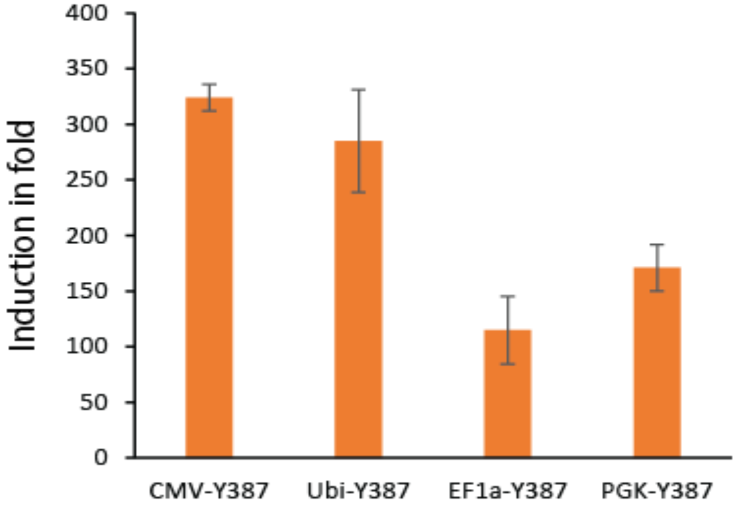


**Fig. S17. The pA regulator functions efficiently with commonly used mammalian promoters.** The Y387 pA regulator coupled to CMV, Ubi, EF1a, and PGK promoter were tested in Hela cells using simple transfection assays and treated with 1μg/mL Tc. Different promoters are expected to have varied transcriptional levels in different cell types, and this may contribute to the varied induction.

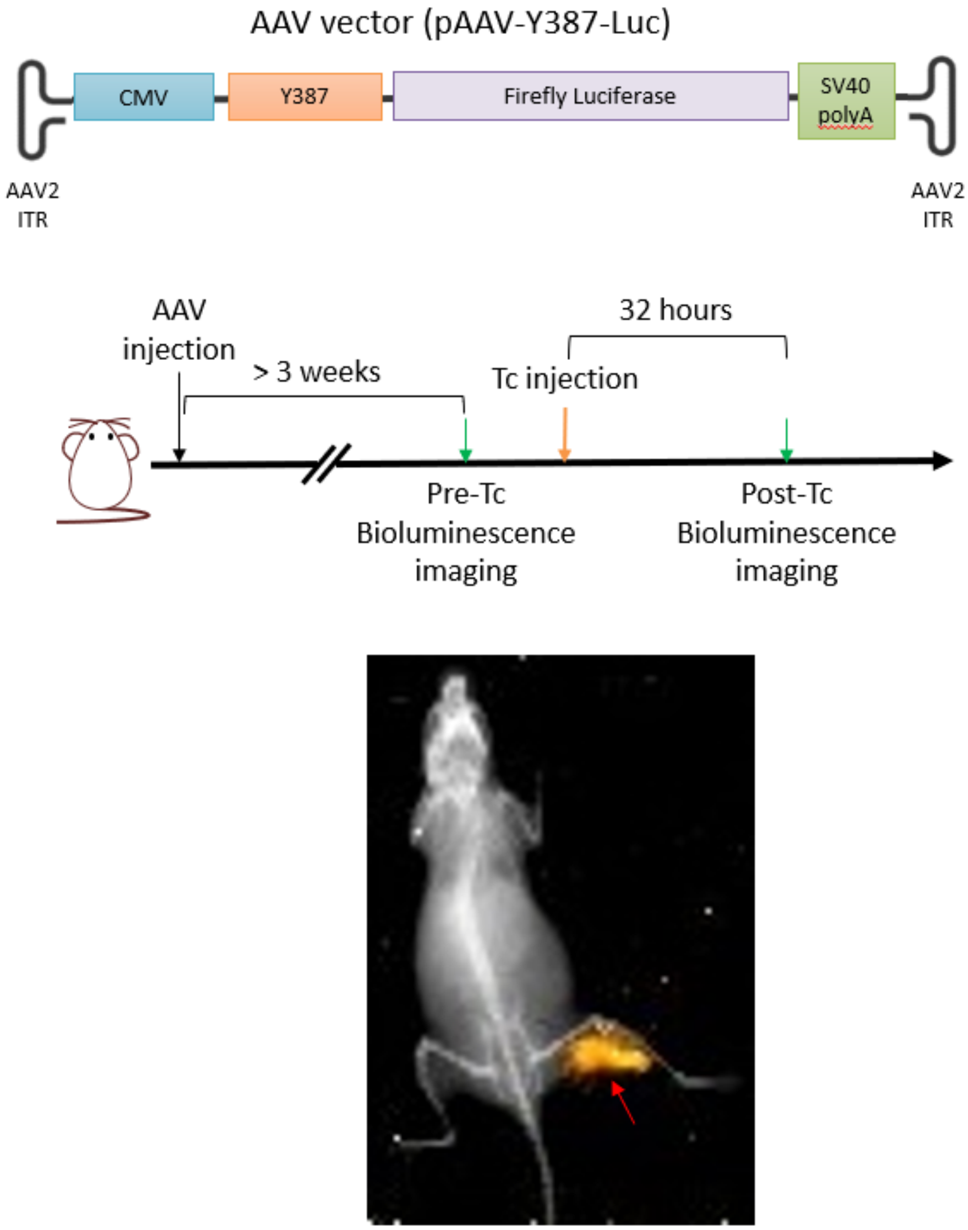


**Fig. S18. Diagram of AAV vector used for *in vivo* study.** Top: The AAV vector encodes a firefly luciferase reporter gene under the control of CMV promoter and Y387 pA regulator. AAV produced for this study has genotype 2 (AAV2) and serotype 9 (AAV9). Middle: Experimental timeline. Female mice of 6 weeks old were used for our study. For each mouse, $6 \times 10^{10}$ AAV particles were injected intramuscularly. Three weeks after AAV injection, mice were imaged for background luciferase expression (Pre-Tc). 32 hours after a single dose Tc injection, mice were imaged again for Tc-induced luciferase expression (Post-Tc). Bottom: X-ray image overlaid with bioluminescence image. Luciferase signal (in yellow) reveals the AAV-injected hind leg muscle. Right hind leg of the mouse was stretched to demonstrate the position of the injected muscle group relative to femur and Tibia-Fibula.
